# Modular Pooled Discovery of Synthetic Knockin Sequences to Program Durable Cell Therapies

**DOI:** 10.1101/2022.07.27.501186

**Authors:** Franziska Blaeschke, Yan Yi Chen, Ryan Apathy, Zhongmei Li, Cody T. Mowery, William A. Nyberg, Angela To, Ruby Yu, Raymund Bueno, Min Cheol Kim, Ralf Schmidt, Daniel B. Goodman, Tobias Feuchtinger, Justin Eyquem, Chun Jimmie Ye, Eric Shifrut, Theodore L. Roth, Alexander Marson

## Abstract

Chronic stimulation can cause T cell dysfunction and limit efficacy of cellular immunotherapies. CRISPR screens have nominated gene targets for engineered T cells, but improved methods are required to compare large numbers of synthetic knockin sequences to reprogram cell functions. Here, we developed Modular Pooled Knockin Screening (ModPoKI), an adaptable platform for modular construction of DNA knockin libraries using barcoded multicistronic adaptors. We built two ModPoKI libraries of 100 transcription factors (TFs) and 129 natural and synthetic surface receptors. Over 20 ModPoKI screens across human TCR and CAR T cells in diverse conditions identified a transcription factor AP4 (TFAP4) construct to enhance long-term T cell fitness and anti-cancer function *in vitro* and *in vivo*. ModPoKI’s modularity allowed us to generate a ∼10,000-member library of TF combinations. Non-viral knockin of a combined BATF-TFAP4 polycistronic construct further enhanced function *in vivo*. ModPoKI facilitates discovery of complex gene constructs to program cellular functions.

**Highlights:**

- Modular pooled knockins of hundreds of TF and surface receptor constructs combined with different antigen receptors
- Chronic stimulation screens discover programs to improve T cell persistence
- Combinatorial knockin screens with ∼10,000 transcription factor combinations
- BATF-TFAP4 dual knockin construct improves CAR T cell function *in vitro* and *in vivo*

## INTRODUCTION

T cells engineered to express transgenic T cell receptors (TCRs) or chimeric antigen receptors (CARs) have emerged as a powerful treatment option for some malignancies by redirecting autologous T cells toward cancer cells (Esensten et al., 2017; Fesnak et al., 2016; June and Sadelain, 2018). While CAR T cells have induced impressive initial response rates in patients with hematologic malignancies, they fail to provide long-term disease-free survival in 40-60% of these patients and have not been successful in most solid tumors (Gardner et al., 2017; Maude et al., 2018). Several factors have challenged adoptive cellular therapies including inadequate antigen targets, cancer escape mechanisms and immunosuppressive tumor microenvironment (Stoiber et al., 2019). In addition to these challenges, T cell function can fail as a result of chronic antigen stimulation or tonic signaling in both TCR- and CAR-based approaches (Delgoffe et al., 2021; Schietinger et al., 2012). Chronically stimulated T cells can differentiate into dysfunctional cell states often characterized by increased cell surface expression of inhibitory receptors (such as PD-1, LAG-3, TIM-3), reduced proliferative capacity and cytokine production, and alterations in transcriptome and chromatin landscapes (Chen et al., 2019; Doering et al., 2012; Man et al., 2017; Martinez et al., 2015; Sen et al., 2016; Wherry and Kurachi, 2015). T cell dysfunction with hallmarks of exhaustion has been identified as a major contributor to poor response to CAR T cell treatment (Fraietta et al., 2018a). Thus, engineering therapeutic T cells with improved fitness in the contexts that otherwise predispose T cells to dysfunction - including chronic antigen stimulation - is a promising strategy to improve clinical responses.

Advances in genome engineering and screening methods have offered numerous approaches to increase fitness of T cell therapies and overcome dysfunctional states. One approach is to tune CAR regulation and signaling itself by either targeting the CAR integration to place it under promoter regulation of the endogenous TCR alpha constant chain (*TRAC*) (Eyquem et al., 2017) or by screening a variety of different co-stimulatory CAR domains to identify CAR designs with favorable phenotypes (Di Roberto et al., 2021; Goodman et al., 2021; Kyung et al., 2021). A second approach uses CRISPR/Cas9 to ablate genes that restrict durable T cell function. CRISPR/Cas9 mediated gene ablation of inhibitory receptors induced in dysfunctional T cells – starting with PD1 - has recently been attempted in clinical trials (Stadtmauer et al., 2020). Loss-of-function screens continue to nominate perturbations that can increase T cell fitness such as knockout (KO) of *Regnase-1* (Wei et al., 2019), *Roquin* (Zhao et al., 2021a), *Ptpn2* (LaFleur et al., 2019), *SOCS1* (Sutra Del Galy et al., 2021) or *RASA2* (Carnevale et al., 2022; Shifrut et al., 2018), in murine and/or human T cells. Third, large-scale gain-of-function screens using either CRISPR activation (CRISPRa) (Schmidt et al., 2022) or a lentiviral library of open reading frames (ORFs) have revealed promising perturbations such as overexpression of lymphotoxin B receptor (LTBR) (Legut et al., 2022). However, these screening approaches were not combined with antigen-specific TCRs or CARs in primary human T cells at scale and CRISPRa screens cannot test synthetic gene products.

One promising approach is to engineer the state of therapeutic TCR or CAR T cells either by direct modulation of transcriptional regulators or through artificial cell surface receptors that alter cellular responses to external cues. For example, overexpression of AP-1/ATF transcription factors (TFs) c-JUN or BATF has been shown to improve CAR T cell function (Lynn et al., 2019; Seo et al., 2021). Numerous groups are now designing synthetic genes encoding “switch” receptors that convert an inhibitory signal into an activating signal by fusing the extracellular domains of inhibitory receptors (e.g. PD-1) to intracellular activating domains (e.g. CD28) (Blaeschke et al., 2021; Liu et al., 2021a; Liu et al., 2016). An array of synthetic surface receptors including CD200R/CD28 and TIM-3/CD28 have been developed (Oda et al., 2017; Zhao et al., 2021b), but systematic analysis is required to learn rules about which extracellular and intracellular domain pairings are most effective. More broadly, a modular screening approach is required to discover specific combinations of TFs or surface receptors (SRs) that can be coupled with specific TCRs or CARs to improve functional performance.

An ideal screening system for multi-gene synthetic programs in therapeutic cells would identify constructs encoding specific TFs or SRs coupled with a specific antigen receptor and targeted to a desired genome location. Targeted pooled CRISPR-mediated knockin screens not only allow for testing of constructs at a specific locus that will be used in ideal therapeutic products, it also overcomes several limitations that have challenged pooled lenti- and retroviral screening approaches: (I) viral recombination (Sack et al., 2016); (II) semi-random integration (Cavazzana-Calvo et al., 2010; Fraietta et al., 2018b); and (III) variable integration copy numbers. We previously developed a non-viral pooled knockin (PoKI) screening platform and screened a 36-member library in combination with an NY-ESO-1-specific TCR after targeted integration into the functionally monoallelic *TRAC* locus of primary human T cells (Roth et al., 2020). However, scaling of this approach was impeded by substantial barcode/construct misassignment due to template switching (∼50%), which limited library size and adaptability.

Here we have overcome these challenges and developed Modular Pooled Knockin (ModPoKI) to rapidly generate and screen functional modules of up to thousands of synthetic sequences (ModPoKI libraries) combined with a specificity module containing a clinically relevant synthetic TCR or CAR sequence. These libraries can be integrated non-virally at targeted genomic sites (Nguyen et al., 2020; Roth et al., 2018), allowing precise control of gene expression and integration number. Barcoded multicistronic adaptors in ModPoKI not only facilitate pooled cloning of modular libraries into knockin constructs, but also can be quantified by simple amplicon sequencing and are compatible with widely used single-cell sequencing workflows. We generated three ModPoKI libraries: a 100-member TF library; a 129-member SR library containing synthetic checkpoint/cytokine/death switch receptors, surface receptors, and chemokine receptors; and a ∼10,000-member combinatorial TFxTF library. The ease of sequencing ModPoKI libraries allowed us to perform over 20 unique pooled screens using these libraries across diverse challenge modules, with each screen replicated in cells from multiple individual human donors to ensure robustness. Using bead stimulation, target-cell stimulation, repetitive stimulation and tonic signaling assays, we identified naturally occurring transcription factors and surface receptors as well as synthetic genes that can improve T cell fitness. Coupling ModPoKI with single-cell transcriptome sequencing (ModPoKI-Seq) revealed transcriptional signatures promoted by promising knockin constructs. The screens nominated synthetic constructs that improve anti-cancer T cell activity *in vitro* and *in vivo,* including a novel TFAP4 and BATF multi-gene knockin. Overall, these studies highlight large-scale modular pooled knockin screens as a powerful method to accelerate synthetic biology programming of cell states with enhanced durability and therapeutic functions.

## RESULTS

### Pooled Non-Viral Knockin Screens Enable Evaluation of Hundreds of Different T Cell Constructs for Cancer Immunotherapy

Recent studies have indicated that reprogramming the transcriptional state of T cells by overexpressing a single transcription factor can confer exhaustion resistance and enhanced therapeutic function (Lynn et al., 2019; Seo et al., 2021). The aim of the first part of our study was to screen 100 transcription factors (TFs) and 129 synthetic and naturally occurring surface receptors (SRs, mostly checkpoint and death switch receptors) in the setting of different TCR/CAR specificities and diverse biological contexts (e.g. single stimulation, repetitive stimulation and tonic signaling) (Figure 1A) to provide a systematic resource of gene constructs that can improve therapeutic T cell functions.

**Figure 1.**
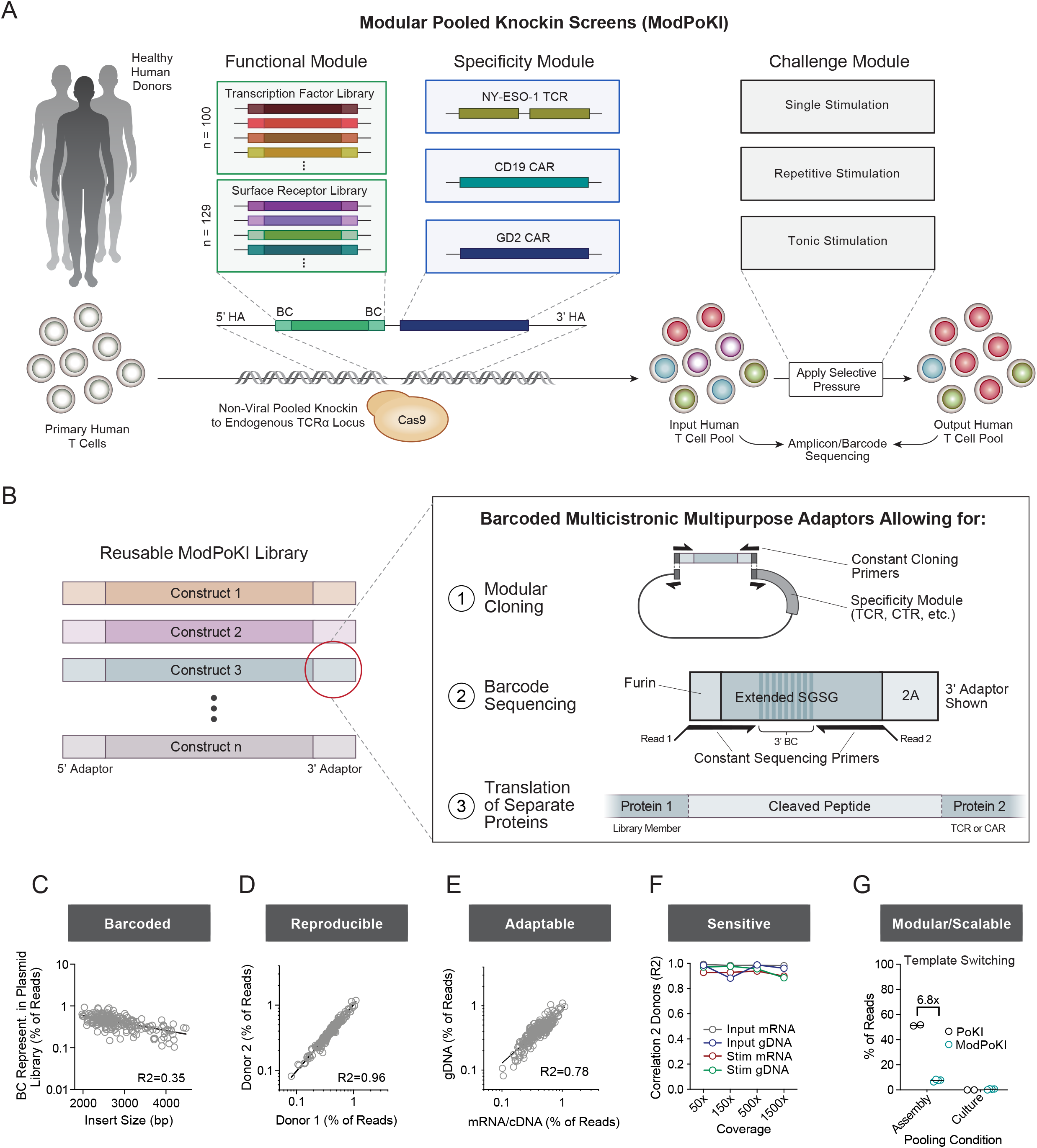
Modular Pooled Knockin of Hundreds of Different T Cell Constructs to Identify Candidates for T Cell Therapy. **(A)** Schematic illustration of the Modular Pooled Knockin (ModPoKI) platform. A surface receptor (SR) or a transcription factor (TF) library (functional module) in combination with an NY-ESO-1 TCR, a CD19 CAR or a GD2 CAR (specificity module) were non-virally integrated into the *TRAC* locus of primary human T cells. The knockin T cell pool was subjected to various screens (challenge module) and construct abundances were analyzed by barcode sequencing. **(B)** Barcoded multicistronic multipurpose adaptors allowed for modular cloning, barcode sequencing and translation of separate proteins. **(C)** Barcode representation in the plasmid library (100 TFs and 129 SRs). N = 2 replicates. Indicated insert size does not include homology arms. R2 was calculated using nonlinear regression (log-log line model, GraphPad Prism). **(D)** Sequencing of the 5’ BC from gDNA after non-viral pooled knockin into cells from primary human donors was reproducible across biological replicates (7 days after electroporation). N = 2 individual donors. R2 was calculated using nonlinear regression (log-log line model, GraphPad Prism). Panel includes data from both the TF and SR library (in combination with the NY-ESO-1 TCR) after KI into primary human T cells. **(E)** Correlation between gDNA and mRNA/cDNA barcode sequencing strategies for one exemplary donor (7 days after electroporation). The second donor confirmed strong correlation (R2=0.76). R2 was calculated using nonlinear regression (log-log line model, GraphPad Prism). Panel includes data from both the TF and SR library (in combination with the NY-ESO-1 TCR) after KI into primary human T cells. **(F)** Donors were highly correlated across different cell coverage ranges, sequencing strategies and experimental conditions (input cells (day 7 after electroporation) vs cells after 4 days of CD3/CD28 bead stimulation (day 11)). N = 2 individual donors. R2 was calculated using nonlinear regression (log-log line model, GraphPad Prism). Panel includes data from both the TF and SR library (in combination with the NY-ESO-1 TCR) after KI into primary human T cells. **(G)** A pilot two-member library of an NY-ESO-1-specific TCR plus GFP vs RFP was pooled at the plasmid assembly stage or after separate electroporation (see Figure S2E). T cells were sorted for TCR knockin and GFP or RFP positivity. Percentage of correctly assigned barcodes was determined by amplicon sequencing (3’ barcode of mRNA/cDNA). The amount of template switching was calculated, extrapolated for an n > 200-member library as described previously (Roth et al., 2020) and compared to the previous version of the pooled KI platform (Roth et al., 2020). Bars represent mean. N = 2 individual donors.

We previously developed a non-viral pooled knockin (PoKI) screening platform to evaluate a 36-member library in combination with an NY-ESO-1-specific TCR (Roth et al., 2020). However, we had observed a significant amount of incorrect barcode/construct assignment due to template switching which prevented pooling of the library at early stages in the protocol and complicated scaling and adaptability of the approach. As we now aimed to screen hundreds to thousands of different T cell constructs and states at the same time in combination with various T cell specificities (CAR or TCR), we developed modular pooled knockin (ModPoKI) screens to minimize template switching and offer a screening platform with exchangeable and adaptable modularity (Figure 1A).

In order to allow highly scalable ModPoKI screens, we generated constructs with multicistronic adaptor sequences that were placed between the DNA sequences of the functional module (TF or SR gene) and the specificity module (CAR or TCR gene) and consisted of barcode-bearing SGSG linkers and cleavage sites (Figure 1B and S1A-C). Each TF or SR library member received one unique barcoded adaptor at the 5’ end and one at the 3’ end to facilitate flexible amplicon sequencing workflows to determine construct identity at either the genomic DNA or mRNA/cDNA level (Figure S1D-F) and enable highly flexible combinatorial approaches. The resulting plasmid library was used to generate double-stranded homology-directed repair (HDR) templates by PCR that were then non-virally integrated into the human TCR alpha constant chain (*TRAC*) locus using CRISPR/Cas9 ribonucleoprotein (RNP) (Figure S2A) (Roth et al., 2018). The modular nature of the approach allowed to combine the TF and SR library with various T cell specificities (CAR or TCR). The resulting CAR or TCR-based T cell pool can be subjected to a variety of different screens and diverse functional assays due to the ease and adaptability of barcode sequencing. The resulting ModPoKI system is barcoded (Figure 1C and S2B), reproducible across donors (Figure 1D and S2C) and adaptable between mRNA/cDNA and gDNA barcode sequencing (Figure 1E and S2D). It is highly sensitive even at low cell coverage (Figure 1F) and modular/scalable due to significantly reduced template switching compared to the previous approach, likely due to improved barcode/insert proximity (Figure 1G and S2E). Pooled knockin single-cell RNA sequencing with barcode sequencing (ModPoKI-Seq) at low coverage confirmed strong correlation of barcode and gene expression (Figure S2F). In summary, ModPoKI screens enable evaluation of hundreds of different T cell constructs for cancer immunotherapy.

### Design of Large Synthetic Libraries for Modular Pooled Knockin Screens

We designed two different knockin libraries to identify constructs that reprogram T cell function through transcription factor overexpression or through altered cell surface receptor signaling. The transcription factor library consisted of 100 members encompassing a variety of different TF families (Figure S3A+B). It contained known regulators of T cell proliferation (such as MYC (Wang et al., 2011)), TFs that have been proposed to increase anti-tumor functions (such as JUN and BATF (Lynn et al., 2019; Seo et al., 2021)), and TFs with unknown functions in the immunotherapeutic context. We covered TFs predominantly expressed in CD4 and CD8 T cells, including TFs that are dynamically regulated upon T cell activation (Figure S3C, https://dice-database.org/). We also included TFs that are naturally predominantly expressed in monocytes, NK cells and B cells to determine if a subset of these could also be used to “synthetically” program improved fitness in T cells (Figure S3C). A list of the tested TFs and their sequences is provided in Supplementary Table 3.

The surface receptor library included mostly synthetic chimeric receptors (also known as “switch receptors”) in which the extracellular domain of an inhibitory checkpoint or death signal receptor was fused to an intracellular domain of an activating receptor to convert inhibitory ligand/receptor interactions into activating signals (Figure S3A and S3D). We used a modular design in which a variety of different extracellular domains were combined with either 4-1BB, CD28, ICOS or other intracellular activation domains. We also included chemokine receptors, cytokine receptors, metabolic receptors, and purely stimulatory molecules into the surface receptor library. A list of the tested SRs and the respective sequences is shown in Supplementary Table 4.

### Discovery of Constructs to Promote Fitness of Stimulated T Cells

We first aimed to identify constructs that could be written into the endogenous *TRAC* locus to enhance fitness of primary human T cells following a single re-stimulation. The NY-ESO-1 TCR in combination with the TF and the SR library was introduced into primary human T cells. The T cell pool was then subjected to various signals including CD3 only stimulation (signal 1 of TCR stimulation through anti-CD3 antibody), CD3/CD28 bead-based stimulation (signal 1 and signal 2 of TCR stimulation), excessive CD3/CD28 stimulation (bead:cell ratio 5:1) and stimulation with NY-ESO-1+ target cells (Figure 2A). To assess effects of targeting both solid tumor and hematological cancer cells, we tested the knockin (KI) pool using two different NY-ESO-1+ target cell lines: 1) A375 melanoma cells that naturally present NY-ESO-1 on HLA-A2, and 2) Nalm-6 leukemia cells transduced to express HLA-A2 with NY-ESO-1. T cells were either stimulated or left resting for four days. RNA was isolated from input cells and from cells on day 4. cDNA was generated and barcode amplicon sequencing was performed to compare the abundance of each construct in the input and output populations. KI of basic leucine zipper (bZIP) TFs, BATF and BATF3, or helix-loop-helix TFs, ID2 and ID3, had the strongest effects on T cell fitness, followed by MYC, a TF known to be essential for T cell proliferation and growth (Wang et al., 2011) (Figure 2B, 2C, S4A and S4B). Among the top negative hits were EOMES, known to be required for full effector differentiation (Pearce et al., 2003) and also associated with exhaustion in anti-tumor T cells (Li et al., 2018), and NFATC1, which has been shown to promote exhaustion in CD8+ T cells (Martinez et al., 2015). Interestingly, BATF or BATF3 KI each seemed to provide an advantage even in the absence of re-stimulation, suggesting potential stimulation-independent effects of these TFs.

**Figure 2.**
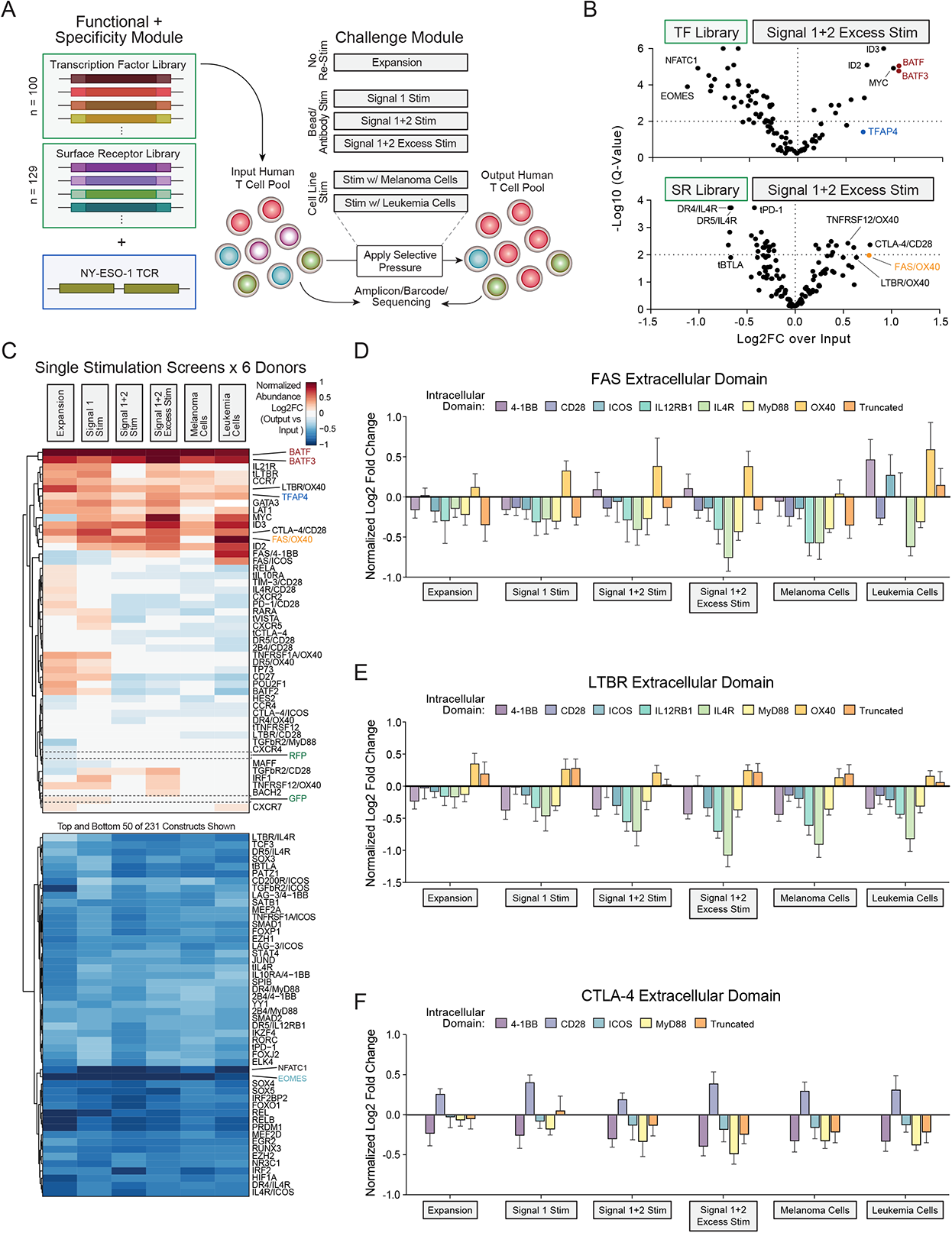
Single Stimulation Pooled Knockin Screens Reveal Known and Previously Undescribed Candidates. **(A)** ModPoKI was performed in primary human T cells with constructs encoding the transcription factor (TF) or surface receptor (SR) library in combination with the NY-ESO-1 TCR. The T cell pool was subjected to multiple different single stimulation screens and knockin cell abundance was assessed by barcode sequencing. Signal 1 Stim = Stimulation with anti-CD3 antibody, Signal 1+2 Stim = Stimulation with CD3/CD28 beads (1:1 bead:cell ratio), Signal 1 + 2 Excess Stim = Stimulation with CD3/CD28 beads (5:1 bead:cell ratio), Melanoma Cells = A375s, Leukemia Cells = Nalm-6 (overexpressing NY-ESO-1). **(B)** Amplicon/barcode sequencing was performed before and after excessive CD3/CD28 stimulation (5:1 bead:cell ratio for four days) to determine log2 fold change in construct abundance in output vs input population. False discovery rate was calculated using the Benjamini-Krieger-Yekutieli method. N = 6 individual donors. **(C)** Representation of T cell constructs was evaluated prior to and after different stimulation conditions. Average log2 fold change over input population is shown (normalized to abundance of RFP and GFP controls). N = 6 individual donors. **(D-F)** Impact of the intracellular domains of the FAS (D), LTBR (E) and CTLA-4 (F) switch receptors was analyzed. Average log2 fold change over input population is shown (normalized to abundance of RFP and GFP controls). N = 6 individual donors. Mean + SEM shown.

Knockin of SR library members could also modulate T cell fitness upon stimulation. Notably, upon excessive stimulation of T cells, death fusion receptors (such as LTBR/OX40, FAS/OX40 and TNFRSF12/OX40) enhanced T cell fitness (Figure 2B, 2C, S4C and S4D). Another hit in this context was the KI construct encoding CTLA-4/CD28, the mouse version of which was shown to increase therapeutic efficacy of donor-lymphocyte infusions (Park et al., 2017; Shin et al., 2012). A novel fusion receptor FAS/OX40 strongly promoted T cell abundance across multiple different screening conditions. Overall, evaluating the fusion receptors across different single stimulation screens, FAS, LTBR, and CTLA-4 extracellular domains tended to perform best (Figure 2C). OX40 intracellular domains performed well with both the FAS and the LTBR extracellular domain (Figure 2D and 2E). Interestingly, the CD28 intracellular domain was the only intracellular domain tested that increased abundance with the CTLA-4 extracellular domain (Figure 2F). These highly parallelized functional assays have potential to inform design of specific fusion receptors that confer context-specific benefits to T cell therapies.

### Repetitive Stimulation Screens Discover TFAP4 KI Improves Antigen-Specific T Cell Persistence

Therapeutic T cells must maintain persistent function through multiple rounds of target recognition if they are to clear large tumor burdens. Unfortunately, repetitive stimulation can lead to T cell dysfunction. To discover constructs that can promote persistent T cell fitness, we performed a repetitive stimulation screen in which we transferred the antigen-specific KI T cell pool to fresh cancer target cells every 48h for five consecutive stimulations (Figure 3A). Pilot experiments with a single knockin construct encoding NY-ESO-1 TCR and a control gene (tNGFR, truncated Nerve Growth Factor Receptor) confirmed that repetitive stimulations with cancer target cells can drive: 1) enrichment of NY-ESO-1 antigen specific cells (Figure S5A), and 2) increasingly differentiated T cell phenotypes of antigen-specific T cells (Figure 3B). Notably, T cells showed increasing expression of exhaustion-associated markers (TOX, LAG-3, TIM-3 and CD39) over the course of the assay (Figure 3C and S5B). RNA-seq confirmed the increased *TOX* expression, along with decreases from peak levels in CD62L (*SELL*), Granzyme B (*GZMB*) and IFN-g (*IFNG*) expression over time, consistent with cellular dysfunction (Figure S5C-E). This *in vitro* model with repetitive exposure to cancer target cells provides an opportunity to discover KI constructs that enhance persistent T cell fitness.

**Figure 3.**
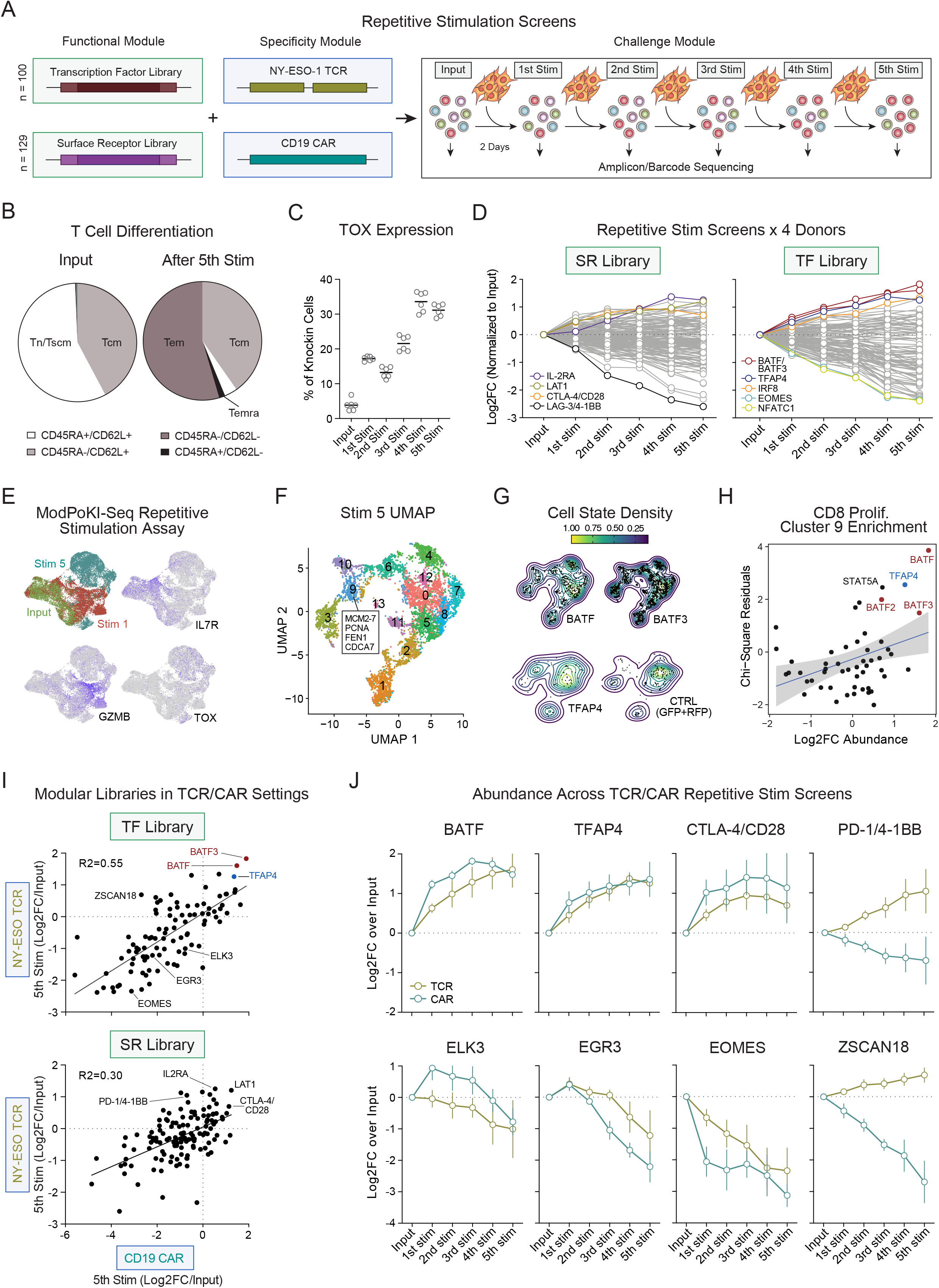
Pooled Non-viral Knockin Screens Identify Highly Functional T Cell Constructs after Repetitive Antigen Encounter. **(A)** Schematic illustration of the repetitive stimulation screen. ModPoKI T cells were generated as described in Figure 1A and stimulated with A375 cells every two days for five consecutive stimulations. Amplicon sequencing was performed at every time point to evaluate abundance of T cell knockins. **(B)** Control T cells (NY-ESO-1 TCR and tNGFR) were generated and subjected to the repetitive stimulation assay to evaluate changes in T cell phenotype. N = 2 individual donors in technical triplicates. **(C)** Single knockin constructs were generated as described in B). Intracellular expression of exhaustion-associated marker TOX was measured by flow cytometry. N = 2 individual donors in technical triplicates. Bars represent mean. **(D)** ModPoKI T cells were generated with constructs encoding the NY-ESO-1 TCR and the transcription factor (TF) or surface receptor (SR) libraries. Average log2 fold change of construct abundance compared to input population is shown. N = 4 individual donors. **(E)** The 100-member TF library was knocked into primary human T cells and single-cell RNA sequencing with barcode sequencing (ModPoKI-Seq) was performed. UMAP plots show overexpression of hallmark genes at the input stage, after one and five stimulations with target cells. N = 2 individual donors. **(F)** Semi-supervised clustering of single cells based on gene expression after five stimulations with target cells. Cluster 9 cells expressed hallmarks of proliferating CD8 cells. Highlighted hallmark genes were derived from top 30 differentially expressed genes. N = 2 individual donors. **(G)** Density plot of top knockin candidates compared to control knockins (GFP+RFP) after five stimulations. N = 2 individual donors. **(H)** Chi-square residuals for enrichment in cluster 9 (proliferating CD8 cells, threshold >30 cells/knockin after 5 stimulations) were compared to abundance log2FC in the bulk screens. N = 2 individual donors for ModPoKI-Seq screen, n = 4 individual donors for bulk abundance screen. **(I)** CD19 CAR TF and SR libraries were generated by pooled assembly. Repetitive stimulation ModPoKI screens were performed and hits in the CAR screens were compared to the TCR screens. N = 4 individual donors for TCR screens, n = 3 individual donors for CAR screens. Nonlinear regression (GraphPad Prism) was used to determine R2. **(J)** Log2 fold changes in abundance were compared between the CAR and the TCR repetitive stimulation screen. N = 4 individual donors for TCR screens, n = 3 individual donors for CAR screens. Mean + SEM shown.

We next introduced the SR or TF library in combination with the NY-ESO-1 TCR into primary human T cells via ModPoKI and used barcode sequencing to monitor abundance of the different constructs throughout each round of the repetitive stimulation assay. Constructs in the SR library encoding the amino acid transporter LAT1 and the high affinity IL-2R (IL2RA) increased in abundance after five target cell stimulations, highlighting that overexpression of natural surface receptors can increase durable fitness in T cells challenged by repetitive stimulation (Figure 3D, left panel, and Figure S5F). In the TF library screen, BATF and BATF3 most strongly promoted T cell fitness over multiple stimulations, consistent with their function in the excessive stimulation screen. In contrast, the EOMES and NFATC1 constructs dropped out suggesting that they limit persistent T cell fitness (Figure 3D, right panel, and Figure S5F). TFAP4 KI emerged as a new hit in the repetitive stimulation assay that was not detected by any of the single stimulation screens. While TFAP4 KI cells increased in abundance following a single stimulation, they did not reach statistical significance as a hit in this context. These results nominated promising constructs and also highlight the importance of testing candidate KI cells in experimental contexts designed to assess persistent T cell fitness.

We next molecularly characterized the effects of TF KIs in the setting of the repetitive stimulation challenge. We coupled ModPoKI with single-cell RNA-seq (ModPoKI-Seq) to discover transcriptomic profiles promoted by each of the 100 TF knockins as T cells were stimulated repetitively. We performed ModPoKI-Seq at the input stage, after one stimulation with target cells (stim 1) and after five stimulations with target cells (stim 5). The input population, stim 1 and stim 5 populations clustered separately with expected expression of hallmark genes identified in each population (Figure 3E). The best performing TF KIs in the fitness screens promoted relatively modest transcriptional changes relative to the transcriptome of control cells (RFP and GFP), while worse-performing constructs often caused a higher variance in gene expression compared to controls (Figure S6A-B). To examine the more subtle beneficial transcriptional changes, we performed semi-supervised clustering of T cell transcriptomes after five stimulations with target cells. This revealed a cluster of CD8 cells characterized by high expression of genes associated with proliferation (Cluster 9), where cells were most strongly enriched for the KIs of top hits in our repetitive stimulation screen including BATF, TFAP4 and BATF3 (Figure 3F-H and S6C). ModPoKI-Seq in the setting of repetitive challenge offers mechanistic insights into gene programs – TFs and downstream target genes – that can be modulated to promote persistent T cell function.

Recognizing the importance of testing the KI genes in therapeutically relevant contexts, we next combined the same TF and SR libraries with a CD19 CAR by pooled assembly (Figure S7A) and assessed whether the same constructs found to be beneficial in NY-ESO-1 TCR+ T cells would also promote fitness of CD19 *TRAC* CAR T cells. Indeed, we observed good correlation of hits for both the TF and the SR library when comparing TCR with CD19 CAR screens (Figure 3I-J and S7B). BATF, BATF3 and TFAP4 constructs all promoted durable fitness with the CD19 CAR in the repetitive stimulation assays, as they had with the NY-ESO-1 TCR (Figure 3I-J). The EOMES KI cells again dropped out with repetitive stimulation of the CD19 CARs (Figure 3J). Interestingly, we identified TFs that had increased abundance after a single stimulation but failed to maintain this advantage after repetitive stimulations (such as EGR3 and ELK3). While many constructs overall performed similarly when combined with a CAR vs a TCR, we observed some constructs (such as PD-1/4-1BB and ZSCAN18) that seemed to have different kinetics in the CAR vs TCR setting (Figure 3J). In summary, repetitive stimulation screens highlighted constructs that may preferentially promote durable fitness through multiple rounds of target cell recognition. The finding that differences in construct performance can occur when paired with a CAR vs a TCR highlights the importance of screening with the exact therapeutic construct that will later be used in the clinic.

### ModPoKI across Dysfunction Screens in TCR and CAR T Cells Confirms Candidate Gene TFAP4

In addition to facing repetitive stimulation as they encounter cancer cell targets, CAR T cells are challenged by variable degrees of tonic signaling, which also has the potential to promote T cell dysfunction (Long et al., 2015). In order to discover synthetic constructs that promote T cell fitness and functionality in the context of a CAR with strong tonic signaling, we cloned our libraries into constructs with the high affinity GD2 CAR (Lynn et al., 2019). Although the high affinity GD2 CAR might drive a less dysfunctional phenotype when placed under *TRAC* promoter control compared to retroviral promoters and non-targeted integrations, we did observe differentiation of GD2 CAR T cells leading to decreased levels of memory markers *SELL* (CD62L), *CCR7, LEF1* and *CD27* and increased levels of exhaustion-associated markers (*TOX, EOMES, LAG3* and *CD244* (2B4)) (Figure S8A). We also observed increased markers of tonic activation in GD2 CAR+ cells compared to bystander T cells (Figure S8B). We next performed pooled knockin of the GD2 CAR in combination with the TF library and compared the performance of the different constructs across dysfunction screens (CD19 CAR or NY-ESO-1 TCR + repetitive stimulation with tumor cells or GD2 CAR tonic signaling) vs single stimulation screens (CD19 CAR or NY-ESO-1 TCR + single stimulation with tumor cells) (Figure 4A). While constructs containing BATF and BATF3 showed increased abundance across multiple screens, TFAP4 overexpressing constructs were more clearly enriched after repetitive stimulation and tonic signaling, suggesting potential benefits in exhaustion-prone environments. TFAP4 was unique among all hits in its strong enrichment trajectory over time in the GD2 CAR screen across four donors (Figure 4B). While only mildly enriched after a single stimulation of CD19 CARs or NY-ESO-1 TCRs, TFAP4 seems to confer strongest advantage in chronic stimulation settings, especially with the tonic signaling GD2 CAR. We next performed arrayed, single non-viral knockins of different CAR constructs in combination with either TFAP4 or a control (tNGFR) for deeper characterization and validation of potential therapeutic benefits conferred by synthetic TFAP4 KI. First, using GD2 CAR T cells, we confirmed that cells with TFAP4 KI constructs expand more than co-cultured tNGFR KI control T cells over time, confirming their competitive fitness advantage (Figure S9A). We next analyzed if TFAP4 KI constructs can improve T cell killing capacity in addition to T cell fitness. We co-cultured GD2+ cancer target cells with GD2 CAR T cells with synthetic TFAP4 KI or with a control KI and indeed observed that the TFAP4 constructs improved cancer killing capacity (Figure 4C, left panel). This effect of TFAP4 KI constructs was confirmed across multiple effector:target (E:T) ratios (Figure S9B). We repeated the experiments with the CD19 CAR, versions of which are already clinically-approved. Notably, CD19 CAR T cells began to demonstrate dysfunctional cancer cell killing *in vitro* after multiple rounds of stimulation (Figure S9C). This dysfunction was mitigated with the TFAP4 KI constructs across multiple E:T ratios (Figure 4C, right panel (with 5x pre-stim), Figure S9D (without pre-stim), Figure S9E and S9F (with 5x pre-stim)). Lastly, recognizing potential safety concerns of increased killing capacity, we confirmed that TFAP4 overexpressing CD19 CAR T cells spare CD19 negative target cells (Figure S9G). In summary, TFAP4 appears to promote persistent and antigen-dependent anti-cancer function.

**Figure 4.**
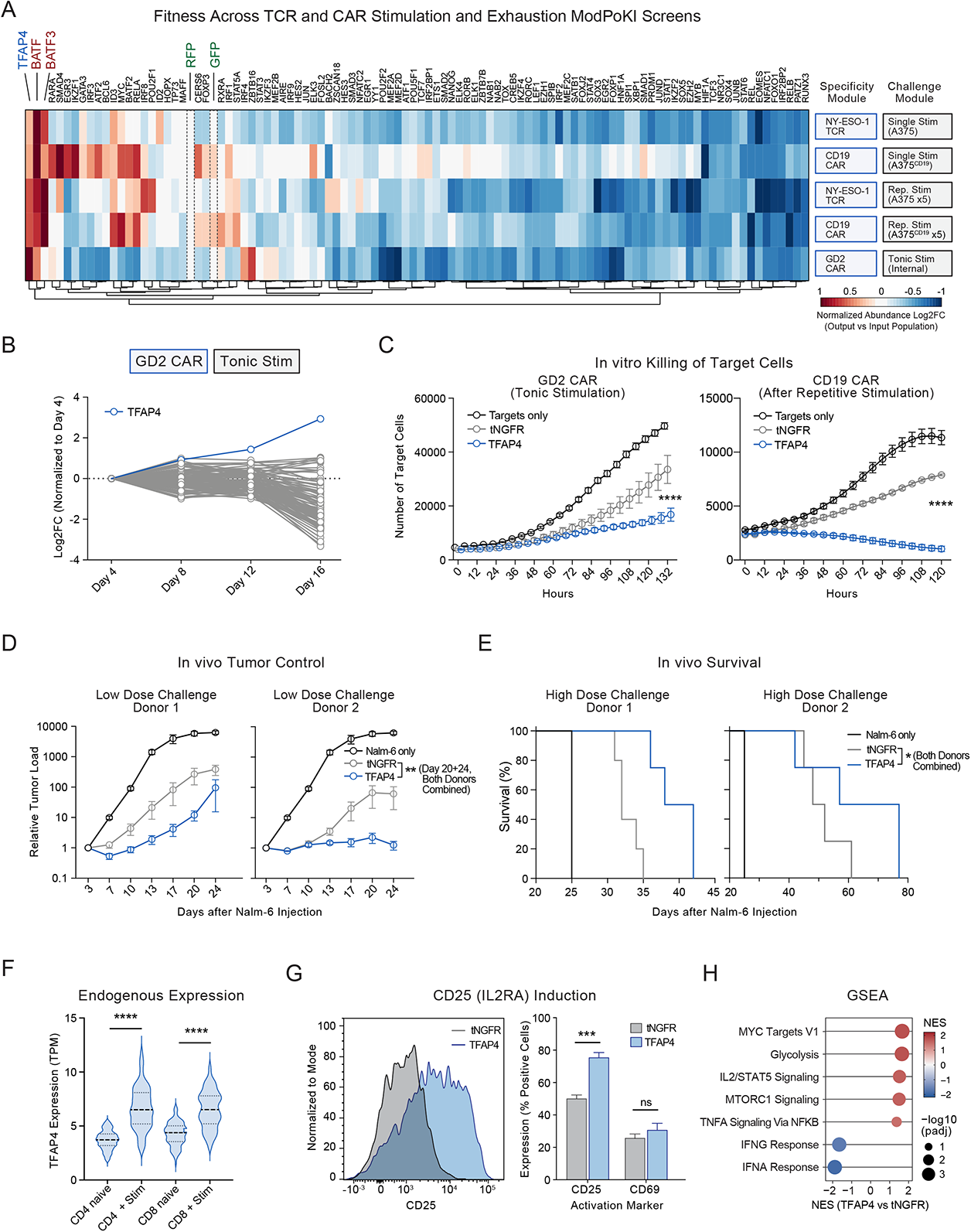
ModPoKI across Dysfunction Screens in TCR and CAR T Cells Nominates Candidate TFAP4 KI. **(A)** ModPoKI screens with the TF library were performed in NY-ESO-1 TCR and CD19 CAR (single vs repetitive stimulation) or GD2 CAR (tonic stimulation) T cells. As the GD2 CAR provides tonic stimulation, GD2 CAR T cells were cultured without addition of target cells. Log2FC in abundance is shown. Heatmap was normalized based on controls (RFP/GFP). N ≥ 3 individual donors per screen. **(B)** Log2 fold changes in the GD2 CAR screen are shown and the strong progressive enrichment of TFAP4 KI cells is highlighted. Mean of n = 4 individual donors. **(C)** Single knockin of the GD2 CAR or CD19 CAR with TFAP4 or control (tNGFR) was performed and cancer cell killing was analyzed by Incucyte measurements. CD19 CARs were prestimulated with target cells five times. N = 2 individual donors per experiment in technical triplicates (GD2 CAR) or quadruplicates (CD19 CAR). Two-way ANOVA test was performed to determine significance including Holm-Sidak’s multiple testing correction. Significance at the last imaging time point is shown; only significance between TFAP4 vs tNGFR is shown, E:T ratio of 1:4 (left panel) and 1:1 (right panel) is shown. Mean + SEM shown. **(D)** NSG mice were challenged with 0.5e6 Nalm-6/GFP/Luc/GD2 cells and treated with 1e6 GD2 CAR+ T cells three days later. Tumor growth was analyzed by bioluminescence imaging. Two individual T cell donors shown (5 mice/donor/construct). Multiple unpaired t test (TFAP4 vs tNGFR) with Holm-Sidak’s multiple comparisons test was performed to determine statistical significance (both donors combined). Mean + SEM shown. **(E)** NSG mice were challenged with 1e6 Nalm-6/GFP/Luc/GD2 cells and treated with 3e6 GD2 CAR+ T cells three days later. Survival analysis for two individual T cell donors shown (≥4 mice/donor/construct). COX regression was performed to determine statistical significance between tNGFR and TFAP4 construct (both donors combined). As the Nalm-6 CAR model is known for outgrowth of antigen-negative (Lynn et al., 2019) as well as antigen-positive tumors that in our experience can occur in body cavities and facilitate endpoint analysis, we used a higher tumor and T cell dose for survival analyses to increase number of mice with clear clinical endpoint signs (hind limb paralysis). **(F)** Endogenous expression of *TFAP4* in naïve vs activated T cells was analyzed using published RNA-seq data (DICE dataset, https://dice-database.org/). Unpaired t test was performed to determine significance. **(G)** CD25 and CD69 expression on GD2 CAR T cells was analyzed on day 8 after electroporation; TFAP4 KI led to increased CD25 surface expression. Multiple t test was performed to determine significance including Holm Sidak’s multiple comparisons test. N = 2 individual donors in technical duplicates. Mean + SEM shown. **(H)** RNA-sequencing of GD2 CAR T cells with TFAP4 or tNGFR KI constructs was performed 7 days after electroporation and showed increased expression of genes belonging to the MYC target family, glycolysis and IL2/STAT5 pathways. N = 2 individual donors.

We next evaluated *in vivo* killing capacity conferred by TFAP4 overexpression in NSG mice that were challenged with GD2+ Nalm-6 leukemia cells (Nalm-6/GFP/Luc/GD2). Anti-GD2 CAR T cells were injected three days after tumor cells and tumor/leukemia growth was analyzed by bioluminescence imaging (BLI) (Figure 4D). TFAP4 KI CAR T cells enhanced leukemia control and increased survival in experiments using T cells from two independent human donors (Figure 4D-E). As the Nalm-6 CAR model is known for outgrowth of antigen-negative (Lynn et al., 2019) as well as antigen-positive tumors that in our experience can occur in body cavities especially after injection of low tumor/T cell numbers and facilitate endpoint analysis, we switched to a higher tumor and T cell dose for survival analyses to increase number of mice with clear clinical endpoint signs due to leukemia progress (hind limb paralysis) in contrast to solid tumor formation in body cavities that is challenging to detect and quantify.

We next evaluated the phenotypic changes synthetic TFAP4 KI induced in primary human T cells. First, we confirmed that non-viral TFAP4 KI can increase TFAP4 expression beyond physiologic levels (Figure S10A-B). TFAP4 is a direct target of MYC expressed after T cell activation (Figure 4F) in an IL-2 dependent manner (Chou et al., 2014) to maintain proliferation (Jung et al., 2008). In line with that, synthetic TFAP4 KI resulted in increased levels of CD25 (*IL2RA*), IL2/STAT5 signaling pathway, MYC target genes, IFN-g, and effector cytokine production, while it decreased IFN-g response genes (Figure 4G-H and S10C-D). Crucially, increases in IFN-g and IL-2 secretion were dependent on the presence of antigen positive target cells (Figure S10E). These results suggest that synthetic TFAP4 KI mediates increased proliferation and antigen-dependent cytokine production, and can promote T cell states with enhanced fitness in the context of chronic antigen challenges.

### Combinatorial Pooled Knockin Screens to Uncover Synergistic Transcription Factor Combinations

TFs can act in combination to reprogram cells to desirable cell states (Hosokawa and Rothenberg, 2021). We wondered if we could discover specific combinations of TFs that work in concert to enhance T cell fitness in the setting of tonic CAR signaling. Analyzing combinations of 100 different transcription factors requires I) knockin library sizes (∼10,000 members) that have not been tested before in this setting and II) successful knockin of very large constructs, especially when done in combination with a CAR (average construct size ∼5.5kb plus homology arms) and thus cannot be performed readily with AAV (adeno-associated virus) HDR templates due to viral packaging size limitations. ModPoKI molecular biology allowed us to overcome these challenges and adapt our ModPoKI screening platform for large-scale combinatorial pooled knockin screens (Figure 5A and S11A). We created a ∼10,000-member library (100 transcription factors plus two controls combined with 100 transcription factors plus two controls) cloned in constructs with the GD2 CAR. Briefly, to accomplish this we first PCR-amplified the GD2 CAR pUC19-based backbone (specificity module) including the homology arms to target the human *TRAC* locus. We next created the two transcription factor inserts by PCR amplification off of the existing TF library using distinct primer pairs for position 1 vs position 2 within the functional module (Figure S11B). The PCRs of the two library inserts were designed to remove the 5’ barcode and constant 5’ linker of the first construct as well as the 3’ barcode and constant 3’ linker of the second construct. By pooled Gibson assembly, a DNA site was created which consisted of the 3’ barcode of the TF in the 1^st^ position, a constant linker (linker 2 – linker 1 junction) and the 5’ barcode of the TF in the 2^nd^ position, creating a unique combinatorial barcode for each TFxTF combination (Supplementary Table 5). HDR templates were generated from the plasmid library by PCR and non-viral knockin of the library into the *TRAC* locus of primary human T cells was performed. Notably, the expected knockin templates spanned a large size range from ∼3.3 to ∼8.2 kb (without homology arms). The fusion region between TF1 and TF2 served as a barcode combination to identify the abundance as well as the orientation (TF1 vs TF2) of the different combinatorial constructs by amplicon sequencing (Figure S11C). Amplicon sequencing of the plasmid pool and the ModPoKI T cell pool 4 days after electroporation confirmed representation of >99% of the constructs at both steps of the protocol, despite the expected construct size-dependent effects on library representation/knockin rate (Figure 5B-C). We were thus able to generate pooled libraries with thousands of different members and successfully achieved diverse knockins including constructs as large as ∼7.6kb based on barcode sequencing.

**Figure 5.**
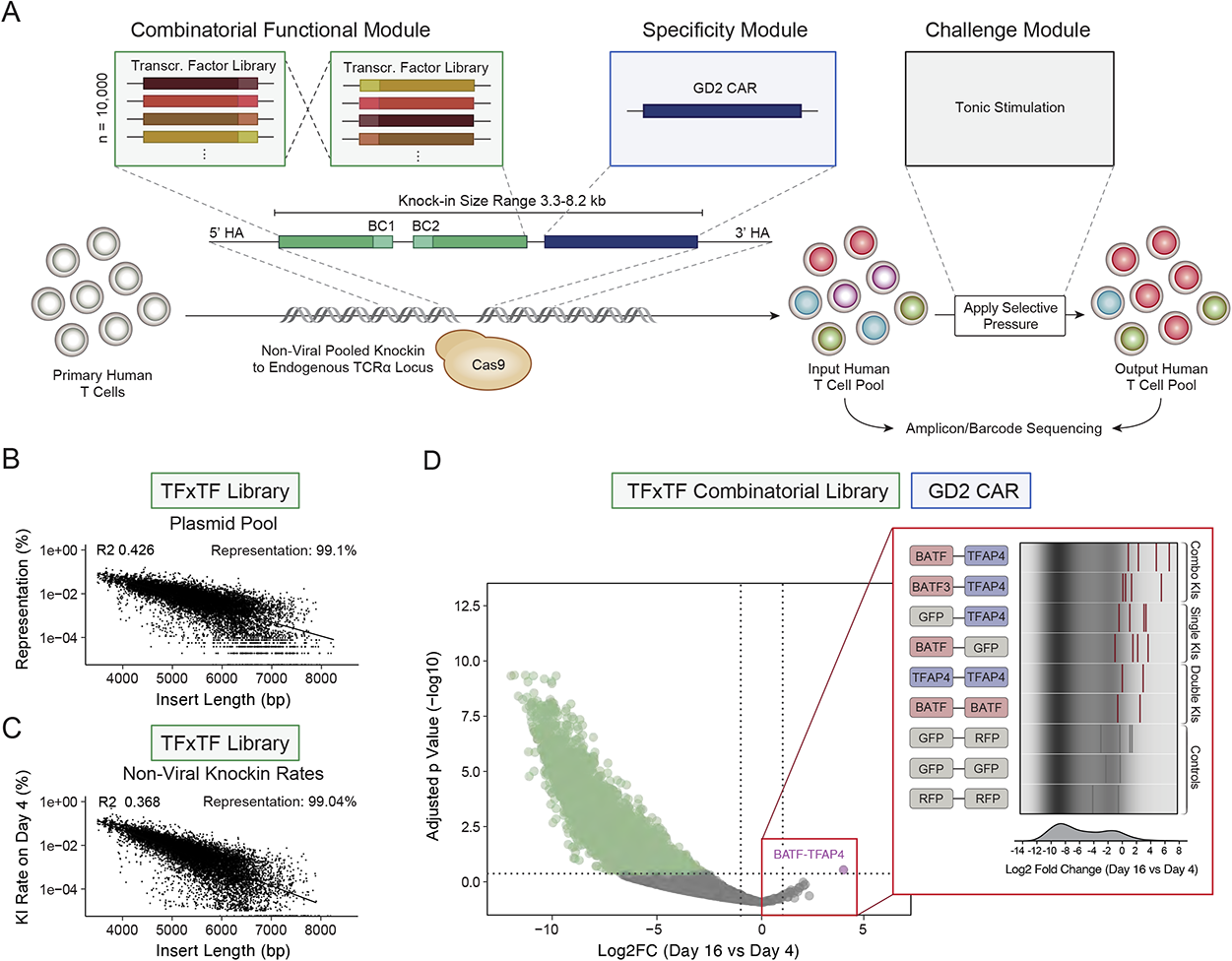
Combinatorial Pooled Knockin Screens Uncover Efficient Transcription Factor Combinations. **(A)** The ModPoKI platform was further advanced to screen pairwise combinations of transcription factors together with the GD2 CAR. In the combinatorial functional module, 100 TFs were combined with 100 TFs (in addition to controls) resulting in ∼10,000 different TF combinations. They were combined with the GD2 CAR resulting in a knockin size range of ∼3.3 to ∼8.2kb. **(B)** Barcode sequencing of the TFxTF combinatorial plasmid library showed size-dependent representation, but confirmed that >99% of constructs were represented after pooled assembly. Statistics were done using linear regression (lm function in R studio). **(C)** Knockin percentage of combinatorial constructs was analyzed in the cell pool on day 4 after electroporation by amplicon sequencing and showed >99% representation of the ∼10,000 constructs. N = 2 individual donors. Statistics were done using linear regression (lm function in R studio). **(D)** The TFxTF combinatorial library was knocked into primary human T cells. As the GD2 CAR provides tonic stimulation, T cells were cultured without addition of target cells. Cells were sorted on day 16 and day 4 after electroporation and the log2 fold change (log2FC) was calculated (day 16/day 4). Log2FC for the combinatorial TFxTF constructs is shown and highlights that combinations of TFAP4 and BATF performed best. N = 2 individual donors. Statistics were calculated using DESeq2. To create the volcano plot, the two possible construct orientations (e.g. BATF-TFAP4 and TFAP4-BATF) were combined to one dataset. The right panel shows barcode representation of the two construct orientations x two donors.

We used combinatorial ModPoKI to test which TFxTF constructs would enhance T cell fitness in the context of tonic CAR signaling. ModPoKI cells expanded in culture due to GD2 CAR tonic signaling. We compared the abundance of each TFxTF combination construct after 16 days in culture to its baseline abundance in the ModPoKI T cell population on day 4 after electroporation. Most TFxTF combinations were depleted from the pool over time, consistent with our previous evidence that major transcriptional changes can be detrimental to fitness (Figure S6A-B). Analysis of the constructs that increased the most in relative abundance (log2 fold change) highlighted that several of the top performing constructs included a combination of TFAP4 and BATF (or BATF3) suggesting that TFAP4 and BATF(3) are key transcription factors that can coordinately drive increased T cell fitness during repetitive simulations (Figure 5D and S11E). Analysis of screens performed in cells from two human donors identified the TFAP4 and BATF combination construct as the most significantly increased in abundance across the two different barcode combinations/knockin directions (TFAP4-BATF and BATF-TFAP4) (Figure 5D). In summary, these data show that large-scale combinatorial knockin screens of ∼10,000 different constructs with an average knockin size of ∼5.5kb is feasible using the ModPoKI screening platform and can help create an atlas of combinatorial KI constructs with potential to enhance therapeutic T cells.

### Combined TFAP4 and BATF KI Induces Favorable States in Therapeutic T Cells

To validate and characterize the benefit of KI constructs combining BATF and TFAP4, we next generated specific knockin constructs with the GD2 CAR and: 1) BATF + TFAP4 combination, 2) single TF + control (RFP-TFAP and BATF-RFP), or 3) control + control (RFP-tNGFR). We performed competitive fitness assays to assess if the combination KI outperformed the individual TF KIs. KI cells with the BATF + TFAP4 combinatorial construct were co-cultured at a ∼50/50 ratio with KI cells with a construct containing only a single TF (along with a control gene), and relative abundance was monitored over time (Figure 6A). KI cells with the BATF + TFAP4 combinatorial construct outcompeted both the TFAP4 + control KI cells and the BATF + control KI cells. The relative benefit of BATF + TFAP4 combination was stronger compared to BATF only than to TFAP4 only, hinting that the majority of fitness benefit (although not all of it) is conferred by TFAP4 KI. Consistent with the effects of the single TFAP4 KI constructs, we found increased levels of CD25 expression in TFAP4-containing combinatorial constructs, whereas CD69 levels were not markedly affected (Figure 6B). When analyzing the phenotype of GD2 CAR T cells 14 days after electroporation, we observed that control (RFP-tNGFR) and BATF KI (BATF-RFP) T cells had high percentages of terminally differentiated Temra cells (CD62L-/CD45RA+), whereas the phenotypes of TFAP4 KI cells (both TFAP4 + control and TFAP4 + BATF KIs) were shifted toward stem cell-like and central memory states with significantly reduced percentages of Temra cells (Figure 6C-D).

**Figure 6.**
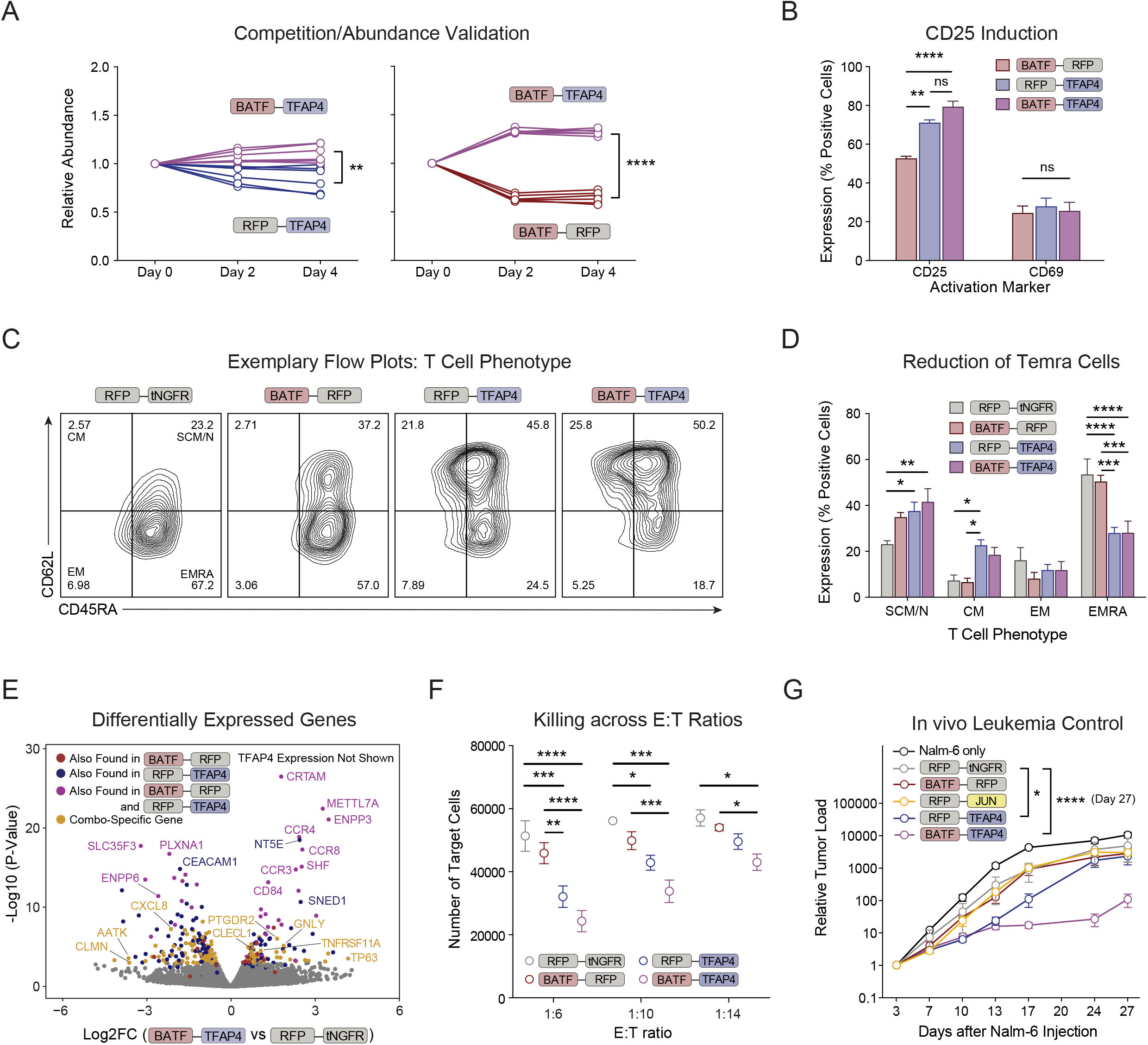
Combinatorial Knockin of TFAP4 and BATF Induces Favorable Transcriptional Programs in Therapeutic T Cells. **(A)** Competitive fitness assays with specific combinatorial knockin constructs (including the GD2 CAR) were performed. ∼50%/50% co-culture analyses of different combinatorial constructs are shown. N = 2 individual donors in technical triplicates. Mean + SEM shown. Unpaired t test on day 4 was performed to determine statistical significance. **(B)** Activation marker expression was analyzed on CAR T cells 8 days after electroporation and showed increased levels of CD25 in TFAP4 KI and BATF-TFAP dual KI CAR T cells compared to BATF KI CAR T cells. N = 2 individual donors in technical duplicates. Mean + SEM shown. 2-way ANOVA with Holm-Sidak’s multiple comparisons correction was performed to determine statistical significance. **(C)** Exemplary flow cytometry plots are shown for phenotypic markers 14 days after electroporation. **(D)** Phenotypic analysis of the different combinatorial KI CAR T cells 14 days after electroporation. N = 2 individual donors in technical duplicates. Mean + SEM shown. 2-way ANOVA with Holm-Sidak’s multiple comparisons correction was performed to determine statistical significance. **(E)** Differentially expressed genes in the BATF-TFAP4 KI CAR T cells compared to RFP-tNGFR control KI CAR T cells were analyzed by RNA-seq 14 days after electroporation, shown in a volcano plot. The most differentially expressed gene was TFAP4 (not shown, log2FC = 5.0, padj = 6.03E-77). The color indicates if the respective gene was also found among the most differentially expressed genes when comparing TFAP4-RFP KI CAR T cells vs control KI CAR T cells, BATF-RFP KI CAR T cells vs control KI CAR T cells or in both of these comparisons. Highlighted in yellow are genes that were differentially expressed selectively in BATF-TFAP4 vs RFP-tNGFR KI CAR T cells. N = 2 individual donors. **(F)** Combinatorial KI CAR T cells were co-cultured with Nalm-6/GFP/Luc/GD2 target cells and target cell killing was analyzed. N = 2 individual donors in technical triplicates. Mean + SEM shown. 2-way ANOVA with Holm-Sidak’s multiple comparisons correction was performed to determine statistical significance. **(G)** To assess *in vivo* function, NSG mice were injected with 0.5e6 Nalm-6/GFP/Luc/GD2 cells IV on day 0 followed by injection of 1e6 GD2 CAR KI T cells IV three days later. Leukemic load was determined by bioluminescence imaging. N = 2 individual T cell donors with 2-5 mice per donor per group. The two different donors are shown separately in Figure S13B. Mean + SEM shown. 2-way ANOVA with Holm Sidak’s multiple comparisons test was performed to compare all constructs against the control (RFP-tNGFR) (both donors combined).

We next evaluated the transcriptional effects of the BATF-TFAP4 combination construct compared to single TF constructs and control constructs. We sorted GD2 CAR+/TCR-cells from each KI population on day 14 after electroporation and performed RNA-seq without addition of target cells (tonic signaling only). Correlation analysis of the log2 fold changes between the respective tested condition and control (RFP-tNGFR) showed that BATF-TFAP4 KI cells were more similar to RFP-TFAP4 KI cells than to the BATF-RFP KI cells (Figure S12A-B). The BATF-TFAP4 KI cells had even less correlation with RFP-JUN KI cells in this setting, suggesting that the transcriptional program promoted by these TFs is divergent from the previously reported program promoted by JUN overexpression (Lynn et al., 2019). Some genes including *CCR3*, *CCR4* and *CCR8* were induced by BATF + TFAP4 KI, BATF-RFP KI and RFP-TFAP4 KI (relative to control KI cells). However, the combined KI of BATF and TFAP4 also promoted differential expression of a variety of genes highlighted in yellow that were not differentially affected by either BATF or TFAP4 KI alone (Figure 6E) such as *TP63* (Tumor Protein 63), *GNLY* (Granulysin), *TNFRSF11A* (Tumor necrosis factor receptor superfamily member 11A) and *CLECL1 (*encoding for the T cell costimulatory protein C-type lectin-like domain family 1). Gene set enrichment analysis of BATF-TFAP4 vs RFP-tNGFR highlighted increased expression of genes involved in cell cycle control (such as E2F targets and G2M checkpoint genes) on day 14 after electroporation, whereas interestingly genes involved in the P53 pathway and IFN-g response seemed to be decreased (Figure S12C, left panel). After stimulation with target cells, BATF-TFAP4 cells had increased expression of genes involved in glycolysis, oxidative phosphorylation, and fatty acid metabolism (S12C, middle and right panels and S12D-E (comparison with single KIs)). Taken together, these results suggest that combinatorial knockin of BATF and TFAP4 can drive both overlapping but also distinct transcriptional changes compared to single BATF or TFAP4 KIs to promote a fitness advantage in the GD2 CAR model of tonic signaling.

The polycistronic TFAP4 single KI construct had improved cancer killing capacity of GD2 *TRAC* CAR T cells *in vitro* and *in vivo*. We now wanted to assess if the TFAP4 and BATF combinatorial KI construct could further enhance the anti-cancer function of GD2 *TRAC* CAR T cells since this combination had conferred an added T cell fitness benefit. Indeed, when co-cultured with GD2 positive cancer target cells, BATF + TFAP4 KI GD2 *TRAC* CAR T cells performed best in an *in vitro* killing assay across multiple E:T ratios (Figure 6F and S13A). Finally, we tested the combinatorial KI cells in the *in vivo* NSG xenograft mouse model of adoptive T cell transfer (Figure 6G and S13B) and observed that KI CAR T cells with the TFAP4 + BATF combinatorial construct showed the best ability to control leukemia growth compared to CAR T cells with either single TF KI construct or with control KI constructs. In summary, combinatorial modular pooled knockin screens of thousands of different synthetic gene constructs can inform design of combinatorial genetic programs that promote enhanced persistence and function to improve adoptive T cell therapy.

## DISCUSSION

Genetically modified T cells are approved for treatment of certain cancer types (Esensten et al., 2017; Fesnak et al., 2016; June and Sadelain, 2018). However, T cell dysfunction resulting from chronic antigen exposure can limit long-term success of adoptive cell therapies (Delgoffe et al., 2021; Schietinger et al., 2012). To discover synthetic knockin constructs that can improve T cell functions, we designed Modular Pooled Knockin (ModPoKI) screening. ModPoKI allowed us to perform a large set of non-viral knockin screens with various antigen receptors and multiple libraries of knockin constructs across various experimental contexts. We designed a 100-member transcription factor and a 129-member surface receptor library that included published switch receptors such as CTLA-4/CD28 and CD200R/CD28 (Oda et al., 2017; Park et al., 2017) and >80 new fusion receptors built by pairing extracellular domains of various cytokine, checkpoint, and death receptors with various intracellular domains of stimulatory receptors. Due to the modularity of this technology, both libraries could be combined with various antigen specificities (CAR or TCR) across multiple individual donors and experimental contexts. Moreover, this advanced screen design allowed for the generation of a combinatorial ∼10,000-member TFxTF library with insert sizes largely exceeding AAV-based knockin capabilities.

Importantly, unlike retro- or lentiviral approaches, ModPoKI uses targeted integration at a defined genomic site. We chose to target the *TRAC* locus as it is functionally monoallelic, knockin can replace the endogenous antigen specificity, the endogenous regulatory elements can drive expression of transgenic CARs and TCRs mimicking expression of endogenous TCRs, and integration of CAR sequences into the *TRAC* locus can reduce the risk of T cell exhaustion (Eyquem et al., 2017; Li et al., 2018; Schober et al., 2019). The targeted integration into the *TRAC* locus instead of semi-random retroviral overexpression could explain why overexpression of the transcription factor c-JUN did not increase durable fitness of tonically signaling GD2 CARs here, in contrast to previous reports (Lynn et al., 2019). Indeed, although we did observe tonic activation and overexpression of hallmark dysfunction genes with the GD2 CAR knocked into the *TRAC* locus, targeted KI of the GD2 CAR may not drive the same extent of dysfunction as the non-targeted viral transduction of the same CAR. Furthermore, expression of c-JUN from a targeted KI driven by the *TRAC* promoter may differ substantially from the levels promoted by viral transduction of the TF and drive a distinct transcriptional program. These results underscore the importance of testing genetic modifications in the same genomic context that will eventually be employed therapeutically in order to identify lead synthetic constructs with the greatest potential to improve immune cell therapies. As cell therapies increasingly rely on targeted modification of genomic loci, ModPoKI is uniquely optimized to compare functional properties of synthetic KI designs.

In order to clear large tumor burdens, therapeutic T cells have to maintain persistent function throughout periods of chronic stimulation from repetitive rounds of antigen encounter or tonic signaling. We thus performed both repetitive stimulation and tonic signaling (GD2 CAR) ModPoKI screens (Lynn et al., 2019). Previous efforts focusing on viral overexpression of bZIP transcription factors have shown enhanced function of GD2, HER2 or CD19 CAR T cells with improved expansion potential, diminished terminal differentiation or enrichment of tumor-infiltrating lymphocytes (Lynn et al., 2019; Seo et al., 2021). Using the ModPoKI platform in combination with repetitive target cell-based CAR or TCR stimulation or tonic signaling, we found that overexpression of the helix-loop-helix transcription factor TFAP4 can promote a proliferative, stem cell-like memory state of therapeutic T cells. Studies in mice have reported that Tfap4 is a Myc-induced transcription factor that maintains Myc-initiated activation and expansion programs in T cells to control microbial infections (Jung et al., 2008). In mice, Tfap4 is regulated by TCR and IL-2R signals and gene-deletion studies indicate that it serves to fine tune clonal expansion of T cells (Chou et al., 2014). Tfap4 has been studied primarily in the context of murine viral infections where it was not essential for short term elimination of the virus, but was crucial in situations where infection could only be controlled by sustained activity of antigen-specific T cells (Chou et al., 2014). These findings align with our discovery that the beneficial effects of the TFAP4 KI constructs are most pronounced in the context of repetitive stimulations or tonic activation of human T cells. Furthermore, Chou et al. described altered glucose utilization in murine Tfap4 knockout cells resulting from defective glycolysis, which could be rescued by retroviral re-expression of Tfap4 (Chou et al., 2014). Here, we report consistent findings that synthetic TFAP4 KI drives increased expression of glycolytic genes. Taken together, TFAP4 KI constructs can drive a transcriptional profile that promotes T cell expansion through chronic stimulation and durable function.

Safety profiles need to be assessed carefully for candidate genetic modifications to promote enhanced expansion and function of cellular therapies. Although we did not observe cytokine release or *in vitro* killing of target cells by TFAP4 KI *TRAC* CAR T cells in the absence of the CAR antigen, safety concerns related to engineered therapeutic cells may eventually warrant the use of suicide switches or synthetic circuits to control expression levels of the transgene (Brandt et al., 2020; Hyrenius-Wittsten et al., 2021; Zhu et al., 2022). Looking forward, ModPoKI could be useful to accelerate the design of these more complex logic-gated synthetic biology programs to enhance cell therapy safety profiles.

Unbiased genome-wide screens now serve as powerful tools to identify candidate genes for gene modification of CAR or TCR T cells. For example, we recently developed a platform to perform genome-wide CRISPR activation (CRISPRa) screens in primary human T cells (Schmidt et al., 2022). However, CRISPRa approaches cannot be immediately translated to the clinic, as they require sustained expression of an activator-linked endonuclease-dead Cas9 (dCas9) which would result in immunogenicity of the therapeutic cells. Nevertheless, genome-wide gain-of-function screens with CRISPRa can be used to nominate genes or pathways that can then be targeted with synthetic knockin constructs and assessed with pooled knockin screens at the appropriate therapeutic locus. For example, both CRISPRa and ORF screens recently nominated overexpression of LTBR as a means to enhance T cell proliferation and cytokine production (Legut et al., 2022; Schmidt et al., 2022). Here, pooled knockin screens revealed LTBR can be engineered into a chimeric receptor (e.g. a LTBR/OX40 fusion protein) that can be knocked into cells along with an antigen receptor to improve fitness. In contrast to CRISPRa screens, pooled knockins allow for screening of both natural and synthetic genes in multicistronic CAR or TCR constructs that can be readily moved toward clinical application without dependence on constant Cas9 expression.

ModPoKI constitutes a platform to rapidly screen hundreds to thousands of different T cell constructs in diverse contexts and across T cell specificities and tumor models. While we have focused on cell proliferation as measured by abundance in this study, pooled knockin screens can be adapted to assess more complex phenotypes such as cytokine release or T cell infiltration into a tumor *in vivo* and will be a crucial tool in the discovery of synthetic genetic modifications that can be engineered to specifically enhance T cell identity and behavior across therapeutic contexts. In the future, modular pooled knockin screens should be readily adaptable to discover constructs that improve function of different CARs or TCRs and even newer synthetic antigen receptors such as HITs (HLA-independent TCRs) (Mansilla-Soto et al., 2022), STARs (synthetic T cell receptor and antigen receptors) (Liu et al., 2021b) or SNIPRs (synthetic intramembrane proteolysis receptors) and SynNotch receptors (Hyrenius-Wittsten et al., 2021; Zhu et al., 2022)). Furthermore, future screens can be performed in regulatory T cells (Tregs) to facilitate the development of treatments for autoimmunity or inflammatory diseases, or in gamma delta T cells. The integration site can be modulated to include loci distinct from the *TRAC* locus, and we anticipate that ModPoKI will be powerful in designing novel gene programs for NK cell therapies, B cell therapies, myeloid cell therapies, iPS cell-derived therapies and beyond. Looking forward, ModPoKI will accelerate candidate selection and design optimization of synthetic biology constructs for basic biological discovery and a diverse array of cellular therapies.

## ACKNOWLEDGMENTS

We thank all members of the Marson Lab; Chris Jeans (QB3 MacroLab) for Cas9 production; Vinh Nguyen (UCSF Flow Cytometry Core, P30 DK063720 and NIH S10 1S10OD021822-01) for his help with sorting; Julia Carnevale, Camillia Azimi, Jacob W. Freimer, Zachary Steinhart, Brian Shy and Sebastian Blaeschke for their helpful suggestions; Stacie Dodgson for critically reading the manuscript and helpful comments. We thank Sarah Pyle for help with illustrations; Jennifer Okano, Jackie Sawin, Jon Woo and Ron Manlapaz for generous assistance. We thank the UCSF Parnassus Flow Core RRID:SCR_018206 and DRC Center Grant NIH P30 DK063720 and NIH S10 1S10OD021822-01 as well as the Gladstone Flow Cytometry Core Facility that was supported by NIH S10 RR028962 and the James B. Pendleton Charitable Trust, DARPA and NIH P30 AI027763. We thank the LARC mouse facilities at UCSF Parnassus Campus and Gladstone Institutes and the sequencing facilities (Functional Genomics Laboratory and Vincent J. Coates Genomics Sequencing Laboratory at UC Berkeley) for generous assistance. We thank Crystal Mackall and Robbie Majzner for providing the GD2 CAR sequence and GD2 expressing cell line (Nalm-6/GFP/Luc/GD2). T.L.R. was supported by the UCSF Medical Scientist Training Program (T32GM007618), the UCSF Endocrinology Training Grant (T32 DK007418), and the NIDDK (F30DK120213). C.T.M. is a UCSF ImmunoX Computational Immunology Fellow and was supported by U.S. National Institutes of Health (NIH) grants F30AI157167, T32DK007418, and T32GM007618. F.B. was supported by the Care-for-Rare Foundation and the German Research Foundation (DFG). F.B. received additional funding for the project through the 2nd Emerging Investigators EHA-EBMT Joint Fellowship Award in the Field of Cell Therapy and Immunotherapy 2022 (sponsored by EBMT and EHA, alongside grants provided by Gilead and Kite). F.B. received the Young Scientists IO research award (sponsored by the Bristol Myers Squibb Foundation) for parts of this work. T.F. was supported by Deutsche Forschungsgemeinschaft (DFG, German Research Foundation) –SFB-TRR 338/1 2021 –452881907. A.M. received funding from the Simons Foundation, Burroughs Wellcome Fund, Career Award for Medical Scientists, The Cancer Research Institute Lloyd J. Old STAR grant, Parker Institute for Cancer Immunotherapy, Innovative Genomics Institute, National Institutes of Health grant P30 DK063720 (Parnassus Flow Cytometry Core, A.M.), National Institutes of Health grant S10 1S10OD021822-01 (Parnassus Flow Cytometry Core, A.M.). A.M. was an investigator at Chan Zuckerberg Biohub and received gifts from the Byers family, Barbara Bakar, Karen Jordan and Elena Radutzky. R.S. was supported by the Austrian Exchange Service- and Austrian Society of Laboratory Medicine as well as the Max Kade Foundation. This work was supported by National Institutes of Health grant S10 RR028962, the Cancer League and the James B. Pendleton Charitable Trust (Gladstone Institutes Flow Cytometry Core Facility).

## AUTHOR CONTRIBUTIONS

F.B., T.L.R. and A.M. designed the study. T.L.R. designed the ModPoKI system. F.B., T.L.R., R.A., R.Y. and Y.C. designed constructs for pooled and single knockins. F.B., Y.C., R.A. and T.L.R. performed pooled knockin experiments. Y.C., R.A., T.L.R., F.B. and E.S. computationally analyzed pooled knockin experiments. F.B., C.T.M. and Y.C. performed bulk and single-cell RNA sequencing experiments. C.T.M., E.S., T.L.R. and F.B. analyzed single-cell RNA sequencing results. C.Y., D.B.G., R.B. and M.K. advised on single-cell RNA analysis. F.B., Y.C., R.A., R.S., W.A.N., A.T. and Z.L. performed *in vitro* and *in vivo* validation experiments. T.F. provided CD19 CAR construct and advised on experiments. J.E., C.Y., E.S. and T.F. advised on the manuscript. F.B., T.L.R. and A.M. wrote the manuscript with input from all authors.

## DECLARATION OF INTERESTS

The authors declare the following competing financial interests: F.B. received the 2nd Emerging Investigators EHA-EBMT Joint Fellowship Award in the Field of Cell Therapy and Immunotherapy 2022 (sponsored by EBMT and EHA, alongside grants provided by Gilead and Kite) and the Young Scientists IO research award (sponsored by the Bristol Myers Squibb Foundation). E.S. served as an advisor for Arsenal Biosciences. J.E. is a compensated co-founder at Mnemo Therapeutics. J.E. is a compensated scientific advisor to Cytovia Therapeutics. J.E owns stocks in Mnemo Therapeutica and Cytovia Therapeutics. J.E. has received a consulting fee from Casdin Capital. The Eyquem lab has received research support from Cytovia Therapeutic and Takeda. T.L.R is a compensated co-founder, is a member of the scientific advisory board, and previously worked as the CSO of Arsenal Biosciences. D.B.G. owns stock in Arsenal Biosciences and served as an advisor for Arsenal Biosciences. D.B.G. is a scientific advisor and stockholder of Manifold Bio, Gordian Biotechnology, Retro Biosciences and NExTNet. C.T.M. is a compensated Bio+Health Venture Fellow at Andreesen Horowitz. R.S. is a scientific advisor of Arsenal Biosciences. A.M. is a compensated cofounder and a member of the scientific advisory boards of Spotlight Therapeutics and Arsenal Biosciences. A.M. serves as member of the board of directors of Spotlight Therapeutics and a board observer at Arsenal Biosciences. A.M. is a cofounder, member of the boards of directors, and a member of the scientific advisory board of Survey Genomics. A.M. is a compensated member of the scientific advisory board of NewLimit. A.M. owns stock in Arsenal Biosciences, Spotlight Therapeutics, NewLimit, Survey Genomics and PACT Pharma. A.M. has received speaking and/or advising fees from 23andMe, PACT Pharma, Juno Therapeutics, Trizell, Vertex, Merck, Amgen, Genentech, AlphaSights, Rupert Case Management, Bernstein and ALDA. A.M. is an investor in and informal advisor to Offline Ventures and a client of EPIQ. The Marson lab has received research support from Juno Therapeutics, Epinomics, Sanofi, GlaxoSmithKline, Gilead, and Anthem. T.L.R., F.B., A.M., R.A., Y.C., C.T.M. and E.S. are listed as inventors on patent applications related to this work.

## LEAD CONTACT

Further information and requests for resources and reagents should be directed and will be fulfilled by the lead contact, Alexander Marson.

## SUPPLEMENTARY FIGURE LEGENDS

**Supplementary Figure S1.**
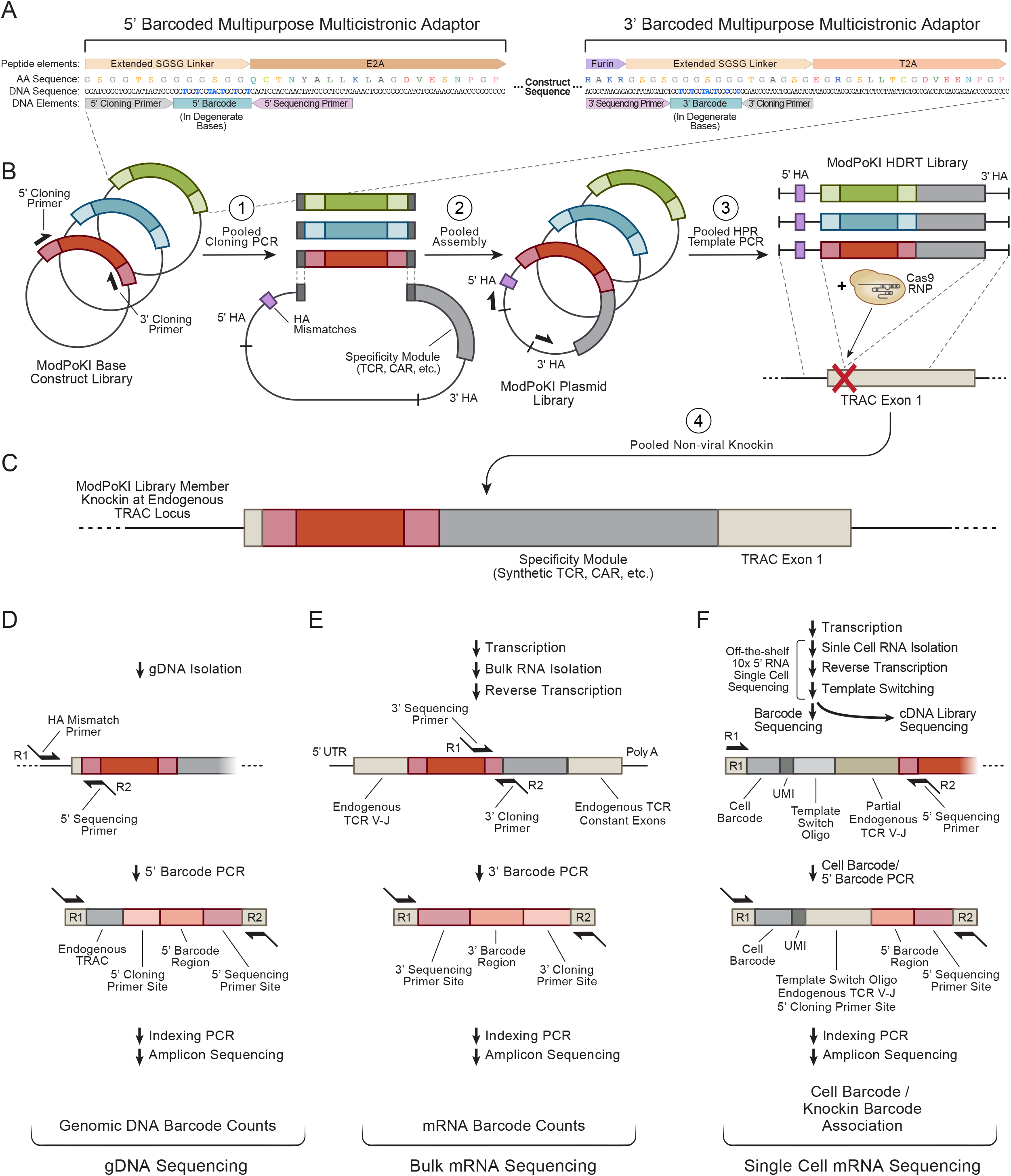
Design of Modular Pooled Knockin (ModPoKI) Constructs, Related to Figure 1. **(A)** To create flexible ModPoKI libraries of sequences that can be integrated into “a functional module” site in pooled knockin templates along with a replaceable antigen receptor in the “specificity module site”, we developed barcoded multicistronic multipurpose adaptors consisting of a 2A site in combination with a DNA barcode. Using degenerate bases, two barcodes per TF/SR member were hidden in the extended SGSG linker of both the 5’ and 3’ adaptor. The adaptors served as binding sites for the constant cloning primers during pooled library assembly and for the constant sequencing primers for barcode readout by amplicon sequencing. **(B)** We synthesized (Twist Bioscience) the initial two ModPoKI libraries, consisting of 100 TFs and 129 SRs plus controls in combination with the NY-ESO-1 TCR. Using the 5’ and 3’ cloning primers (ModPoKI_Cloning_Insert_forw/rev), the inserts were amplified in a pooled PCR reaction and then integrated by pooled Gibson assembly into the final plasmid backbone that contained the specificity module (e.g. CD19 or GD2 CAR) as well as the homology arms. The 5’ homology arm contained DNA mismatches to allow amplicon sequencing off the gDNA without sequencing the dsDNA template. HDR templates were generated by pooled PCR resulting in double-stranded ModPoKI HDRT (homology-directed repair template) libraries. **(C)** The ModPoKI HDRT libraries were used for pooled non-viral knockin into exon 1 of the *TRAC* locus by ribonucleoprotein (RNP) electroporation of Cas9, a *TRAC*-targeting gRNA complex, and the templates. The readout of the screens was done by either sequencing of gDNA (D), mRNA/cDNA (E) or using commercially available scRNA reagents (F). **(D)** Genomic DNA was isolated from primary human T cells after targeted knockin of the ModPoKI library. The barcode region was amplified and Illumina Read1 and Read2 sequences were added using the reverse 5’ sequencing primer in combination with the homology arm mismatch forward primer (ModPoKI_5’_DNA_forw/rev) to prevent sequencing of the template instead of the knocked in DNA sequence. After a subsequent indexing PCR, amplicon sequencing was performed to analyze genomic DNA barcode counts. **(E)** mRNA was isolated from primary human T cells after targeted knockin of the ModPoKI library and reverse transcribed into cDNA. The barcode region was amplified and Illumina Read1 and Read2 sequences were added using the reverse 3’ sequencing primer in combination with the forward 3’ sequencing primer (ModPoKI_3’_RNA_forw/rev). After a subsequent indexing PCR, amplicon sequencing was performed to analyze mRNA/cDNA barcode counts. **(F)** Using off-the-shelf 10x 5’RNA single-cell sequencing technology, GEMs (Gel Bead-In Emulsions) were generated and barcoded followed by reverse transcription, template switch oligo priming and transcript extension. cDNA was amplified, quantified and then split into separate fractions for barcode sequencing (25%) and single-cell transcriptomes (75%) (Roth et al., 2020). All primer sequences are listed in Supplementary Table 1.

**Supplementary Figure S2.**
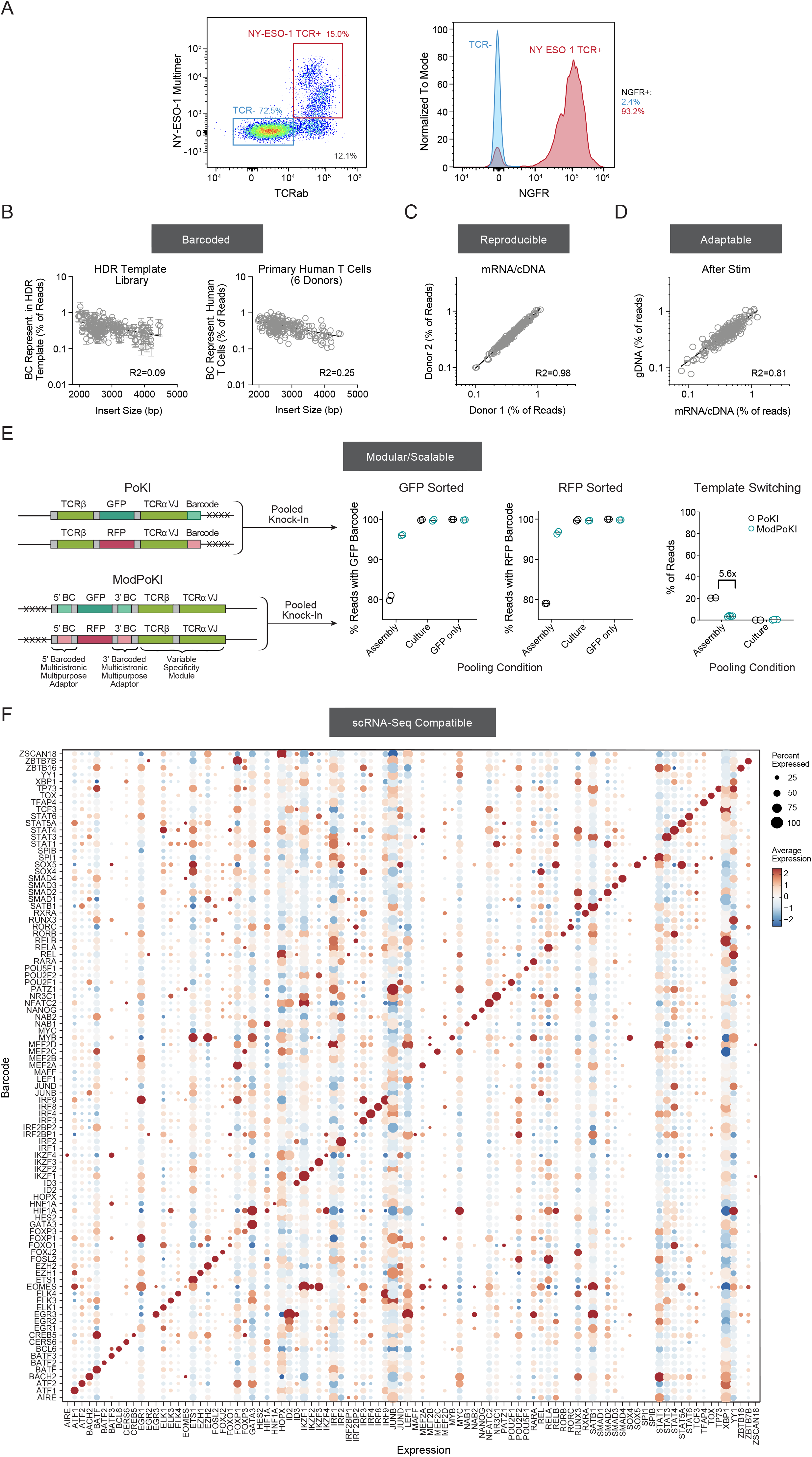
Quality Control Metrics of Non-viral Knockin and Pooled Knockin Libraries, Related to Figure 1. **(A)** Exemplary flow cytometry assessment after knockin of an NY-ESO-1 TCR tNGFR construct including the new barcoded multicistronic adaptor sequences into primary human T cells. **(B)** A mild negative correlation between construct size and library representation was observed in the HDR template pool (n = 4) and of knockin reads in six human donors (day 7 after electroporation). Insert size refers to the number of base pairs that gets inserted into the genomic DNA (without the constructs’ homology arms). R2 was calculated using nonlinear regression (log-log line model, GraphPad Prism). 100 TF and 129 SR library members shown. **(C)** Sequencing of the 3’ BC from mRNA/cDNA was highly reproducible across biological replicates (day 7 after electroporation). N = 2 individual donors, one per axis. R2 was calculated using nonlinear regression (log-log line model, GraphPad Prism). **(D)** Correlation between gDNA and mRNA/cDNA barcode (BC) sequencing strategies. 5’ BCs sequenced off gDNA and 3’ BCs sequenced off mRNA/cDNA from the same pooled knockin experimental donor were well correlated after stimulation with CD3/CD28 beads for 4 days (day 7-11 after electroporation). One exemplary donor shown. Results from a second donor were also correlated (R2=0.66). R2 was calculated using nonlinear regression (log-log line model, GraphPad Prism). TF and SR libraries shown. **(E)** A pilot library of an NY-ESO-1-specific TCR plus GFP vs RFP was pooled at the plasmid assembly stage or after separate electroporation. T cells were sorted for TCR knockin and GFP or RFP positivity and percentage of correctly assigned barcodes was determined by amplicon sequencing (3’ barcode was sequenced off of cDNA). Percent reads with correctly assigned barcodes in sorted populations was notably improved in the new version (ModPoKI) over PoKI v1 (Roth et al., 2020) when pooling at the assembly state. Amount of template switching was calculated for the n = 2-member library and compared to the previous version of the pooled KI platform (Roth et al., 2020). The new screening platform led to 5.6x decreased observed template switching in the n = 2-member library (extrapolated data for the n > 200-member library shown in Figure 1G). Bars represent mean. N = 2 individual donors. **(F)** The transcription factor library in combination with the NY-ESO-1 TCR specificity module was knocked into primary human T cells. Cells were sorted for NY-ESO-1 TCR expression and ModPoKI-Seq (barcode sequencing and transcriptome sequencing) was performed. Cell identity (based on barcode sequencing) was well correlated with transcript expression of the knocked in TF confirming successful knockin and overexpression on RNA level. TF constructs that were codon-optimized or showed no barcode/transcript expression were removed from the analysis. N = 2 individual donors.

**Supplementary Figure S3.**
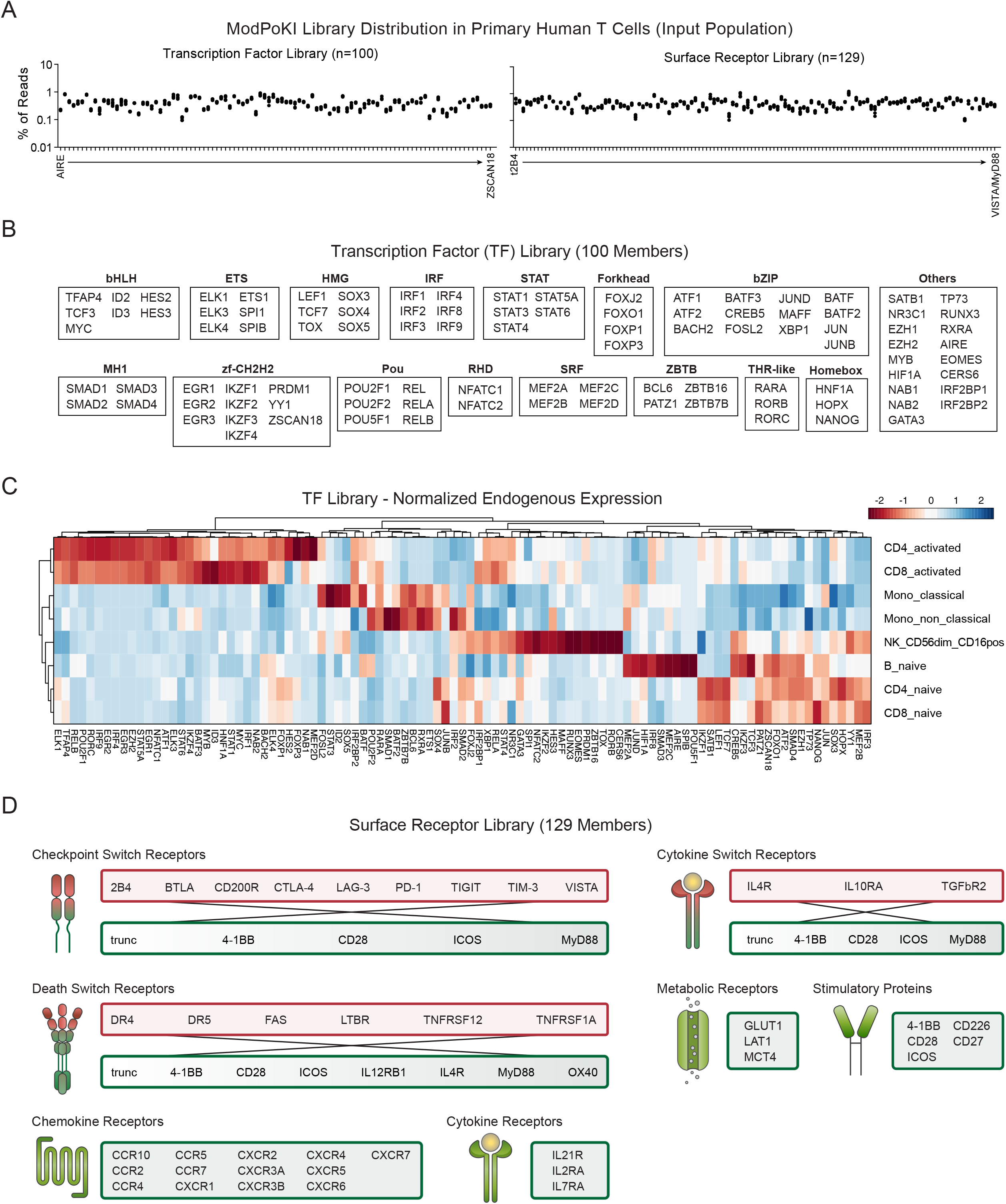
Characterization of Pooled Knockin Libraries, Related to Figure 1. **(A)** Percent of total reads of pooled knockin libraries in six human donors. Transcription factor (TF) and surface receptor (SR) libraries were knocked in as one large library and subsequently computationally separated into individual libraries for analysis. Construct barcodes were consistently well-represented with even library distribution. **(B)** Schematic illustration of the transcription factor library (n = 100) and assignment to the respective transcription factor families. Construct list and sequences can be found in Supplementary Table 3. **(C)** Endogenous expression of the transcription factor library members based on published RNA-seq data (DICE dataset, https://dice-database.org/). Expression was scaled by column. SOX3 was excluded from the heatmap since it had no expression in the tested cell types according to the DICE database. **(D)** Schematic illustration of the surface receptor library (n = 129) and assignment to different receptor groups. Construct list and sequences can be found in Supplementary Table 4.

**Supplementary Figure S4.**
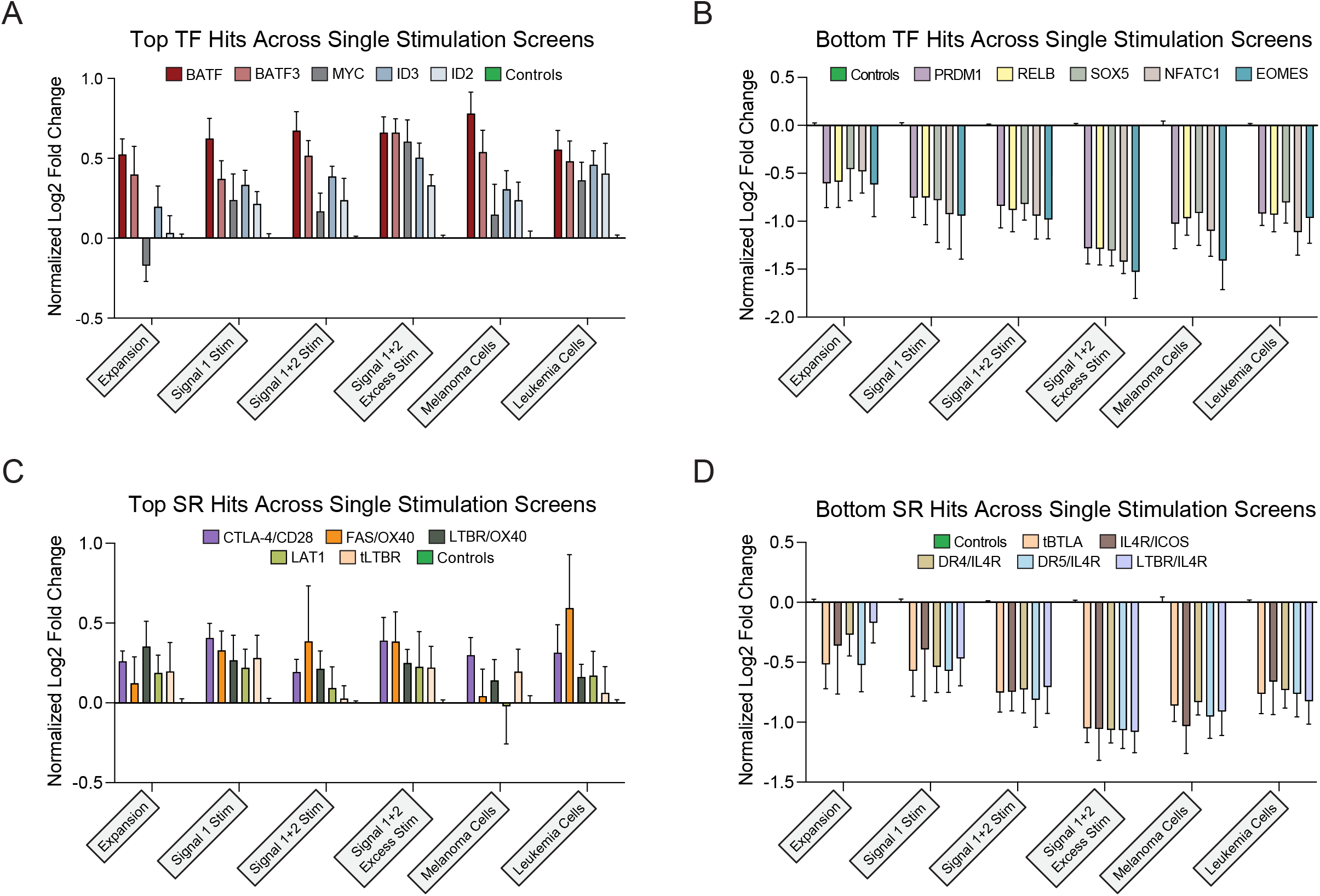
Single Stimulation Pooled Knockin Screens Reveal Known and Previously Undescribed Candidates, Related to Figure 2. **(A-D)** Pooled knockin with constructs encoding the transcription factor (TF) or surface receptor (SR) library in combination with the NY-ESO-1 TCR was performed in primary human T cells. Resulting cells were subjected to the screens described in Figure 2. Log2 fold change in abundance for top and bottom hits (based on excessive stimulation) across single stimulation screens is shown (normalized to abundance of GFP and RFP controls). N = 6 individual donors. Mean + SEM shown.

**Supplementary Figure S5.**
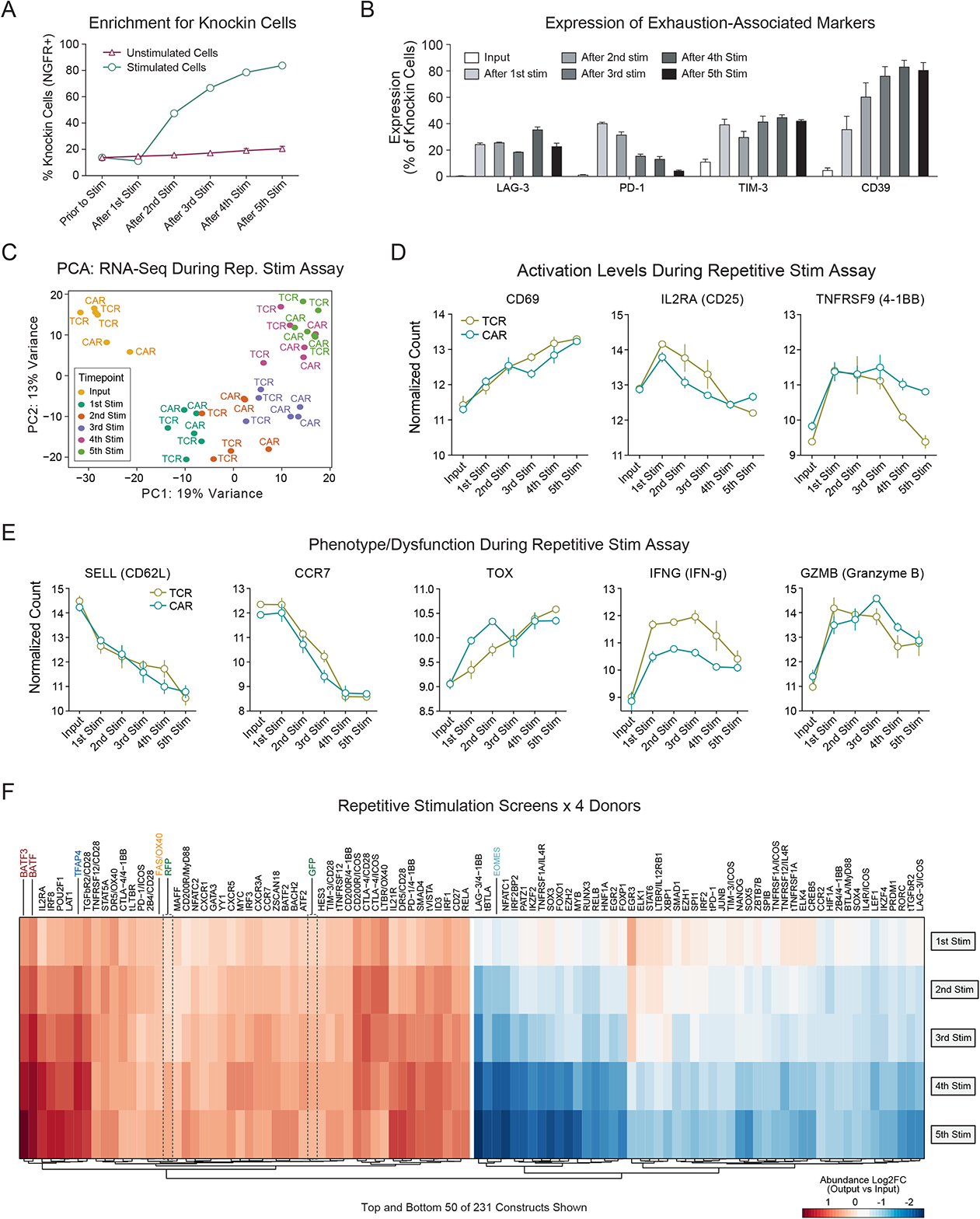
Repetitive Stimulation Assay Setup and Screening Data, Related to Figure 3. **(A)** Control T cells (NY-ESO-1 TCR plus tNGFR) were subjected to the repetitive stimulation screen described in Figure 3A. Knockin percentage (NGFR+) was determined by flow cytometry during the course of the assay and compared to T cells without re-stimulation. N = 2 individual donors in technical triplicates. Mean + SEM is shown. **(B)** Surface expression of exhaustion-associated inhibitory molecules LAG-3, PD-1, TIM-3 and CD39 was analyzed by flow cytometry through the course of the assay. N = 2 individual donors in technical triplicates. Mean + SEM shown. **(C)** Control T cells (NY-ESO-1 TCR or CD19 CAR plus tNGFR) were generated and bulk RNA-seq analysis was performed at every time point of the repetitive stimulation assay as previously reported in a manuscript accepted for publication ((Carnevale et al., 2022), GSE204862). Principal component analysis is shown. N = 3 individual donors. **(D)** Bulk RNA-seq ((Carnevale et al., 2022), GSE204862) revealed progressive induction of *CD69* during the repetitive stimulation assay while *IL2RA* and *TNFRSF9* expression peaked relatively early and then decreased. N = 3 individual donors. Mean + SEM shown. **(E)** A variety of other markers of T cell phenotype, exhaustion and effector function were analyzed and compared between CAR and TCR control T cells ((Carnevale et al., 2022), GSE204862). N = 3 individual donors. Mean + SEM shown. **(F)** The TF and SR libraries were knocked into primary human T cells in combination with the NY-ESO-1 TCR and subjected to the repetitive stimulation assay. Mean log2FC (output vs input) is shown. N = 4 individual donors. Top and bottom 50 of 231 constructs are shown.

**Supplementary Figure S6.**
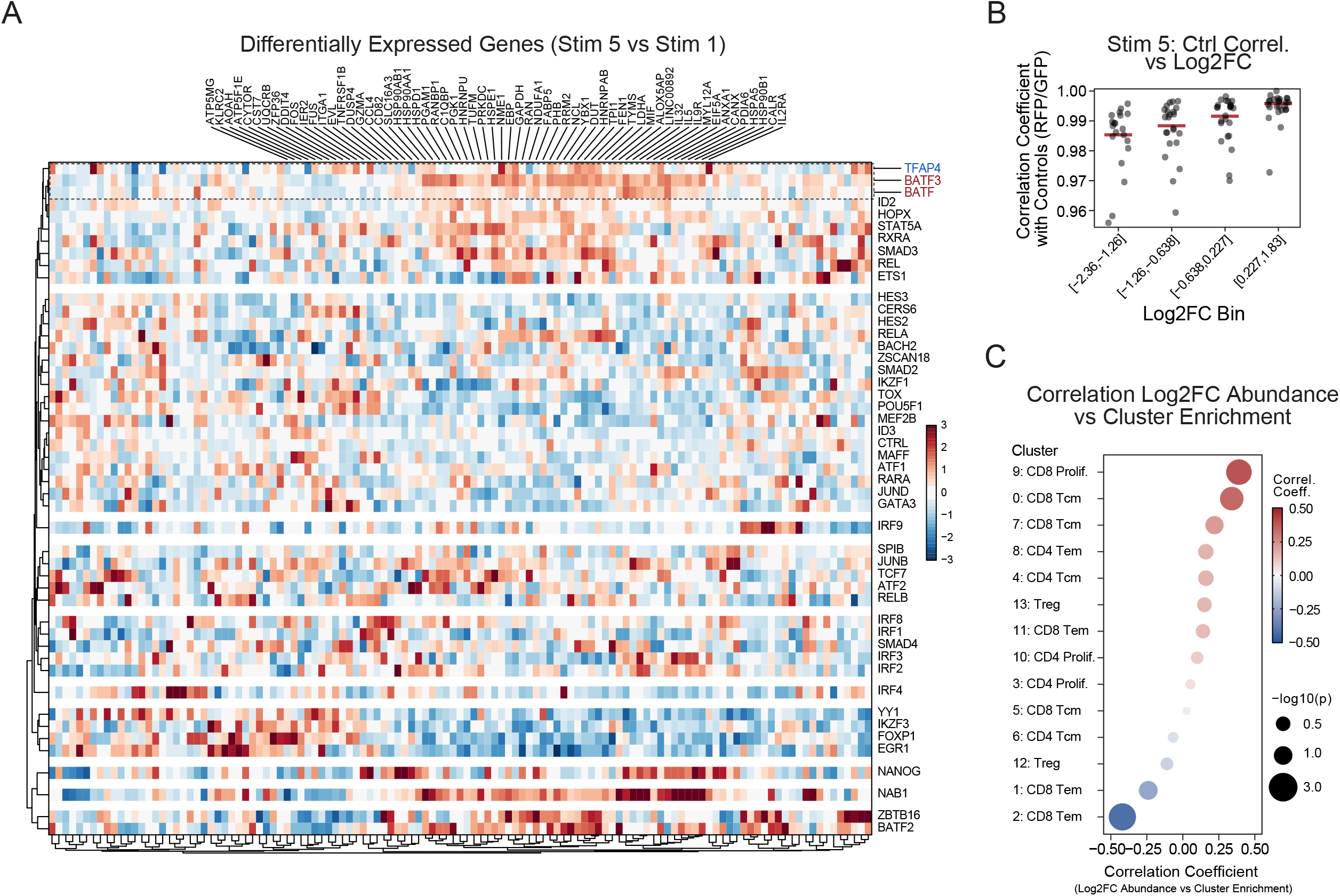
Repetitive Stimulation ModPoKI-Seq Screening Data, Related to Figure 3. **(A)** The 100-member TF library was knocked into primary human T cells from n = 2 individual donors in combination with the NY-ESO-1 TCR. ModPoKI-Seq (single-cell transcriptome coupled with KI barcode sequencing) was performed. Heatmap shows differentially expressed genes (stim 5 vs stim 1) of the different knockins (threshold >30 cells per knockin after 5 stimulations). **(B)** Log2FC bins were generated based on abundance log2FC in the bulk NY-ESO-1 TCR repetitive stimulation fitness screen and compared to the correlation in gene expression with controls (RFP/GFP) in ModPoKI-Seq. Best-performing knockins in fitness screens showed highest correlation coefficients with controls. N = 2 individual donors for ModPoKI-Seq screen, n = 4 individual donors for bulk fitness screen. **(C)** Correlation between cluster enrichment in ModPoKI-Seq (threshold >30 cells per knockin after 5 stimulations) and abundance log2FC in bulk repetitive TCR stimulation fitness screens revealed highest correlation score for enrichment in the CD8 proliferating cluster 9. N = 2 individual donors for ModPoKI-Seq screen, n = 4 individual donors for bulk fitness screen.

**Supplementary Figure S7.**
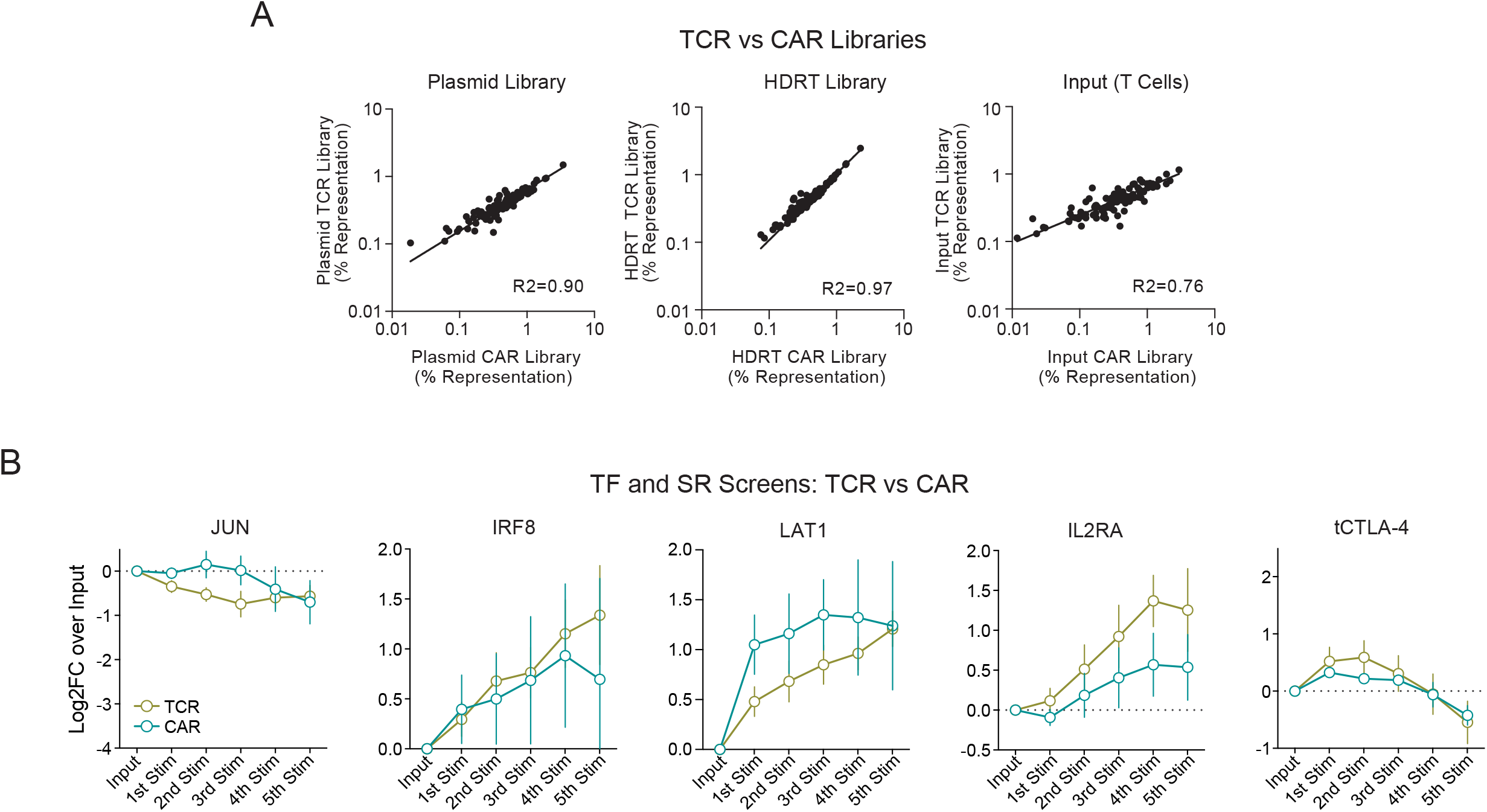
Construction of CD19 CAR Libraries and Construct Performance in Repetitive Stimulation Screens, Related to Figure 3. **(A)** To generate the CD19 CAR library, a CAR plasmid containing the *TRAC*-derived homology arms, the CD19 CAR (FMC63) as well as constant linkers was generated and linearized by PCR. The TFs/SRs plus constant linkers were amplified from the TCR library by PCR. The CAR backbone plus TF/SR inserts were linked using pooled Gibson assembly. Representation of different SR and TF library constructs was analyzed by amplicon sequencing of the plasmid pools, the HDR template libraries and the T cell pool seven days after non-viral knockin (input population for the screens). N = 1 for CAR plasmid, n = 2 for TCR plasmid, n ≥ 3 for HDR templates and input population (individual donors). R2 was calculated using nonlinear regression (log-log line model, GraphPad Prism). **(B)** Log2 fold changes in abundance were compared between the CAR and the TCR repetitive stimulation screen and showed comparable trends for most constructs. While IRF8, LAT1 and IL2RA increased in abundance over time, JUN did not show a significant increase in abundance. Interestingly, control construct tCTLA-4 trended to increase after initial stimulations but dropped out later in the assay (in contrast to e.g. CTLA-4/CD28 fusion, see Figure 3J). N = 4 individual donors for TCR screens, n = 3 individual donors for CD19 CAR screens. Mean + SEM shown.

**Supplementary Figure S8.**
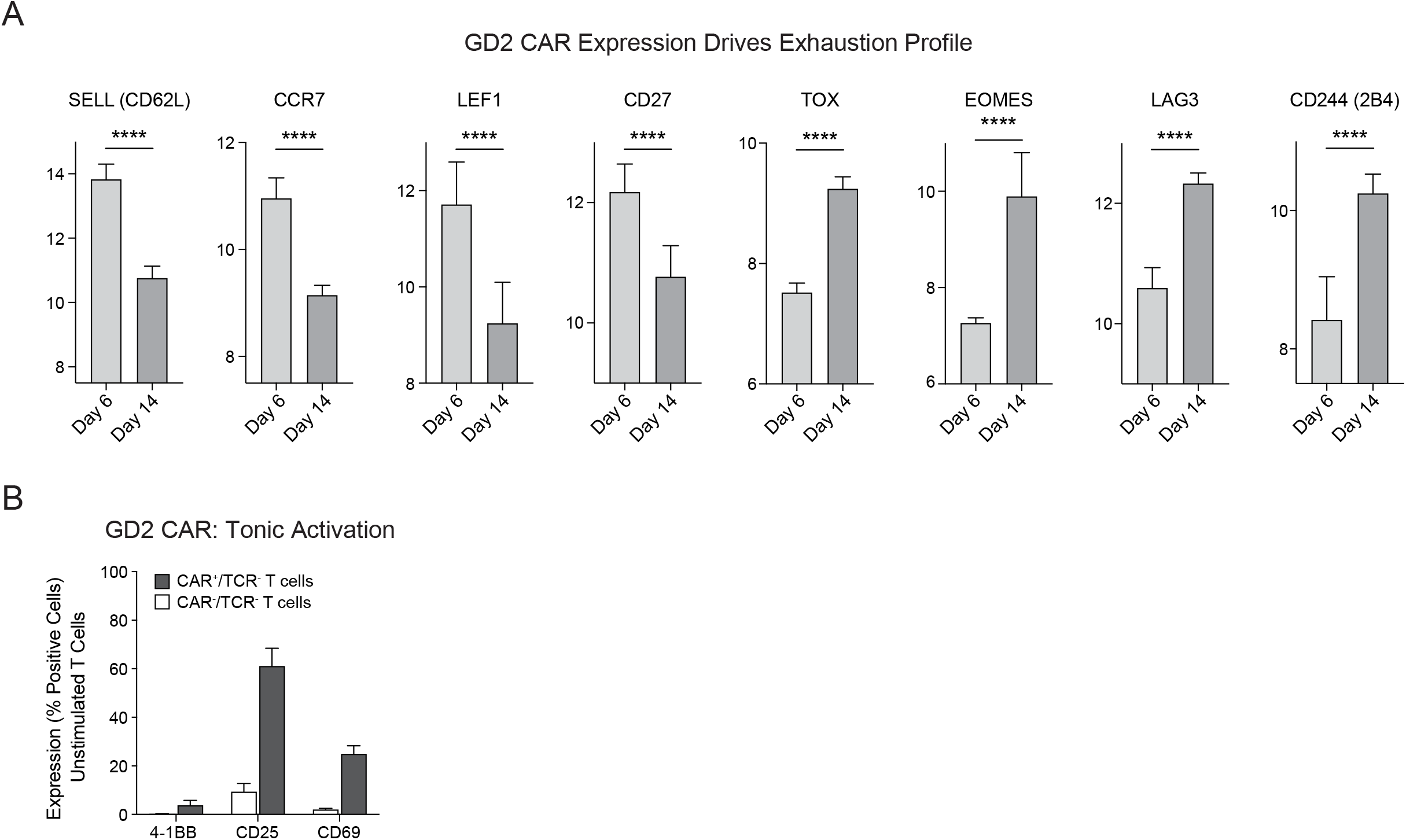
Characteristics of the GD2 CAR Under *TRAC* Promoter Control, Related to Figure 4. **(A)** RNA-seq was performed on GD2 CAR T cells 6 days and 14 days after electroporation. On day 14, GD2 CAR T cells showed decreased levels of early T cell differentiation/memory markers *SELL*, *CCR7*, *LEF1* and *CD27* and increased levels of dysfunction markers *TOX*, *EOMES*, *LAG3* and *CD244*. N = 2 individual donors. Statistical significance was analyzed using DESeq2. Mean + SEM shown. **(B)** Flow cytometric analysis of GD2 CAR+ vs bystander CAR-cells reveal elevated expression of activation markers CD25 and CD69 on CAR T cells even in the absence of target cells 8 days after electroporation, consistent with tonic CAR signaling similar to what was previously described (Lynn et al., 2019). N = 2 individual donors in technical triplicates. Mean + SEM shown.

**Supplementary Figure S9.**
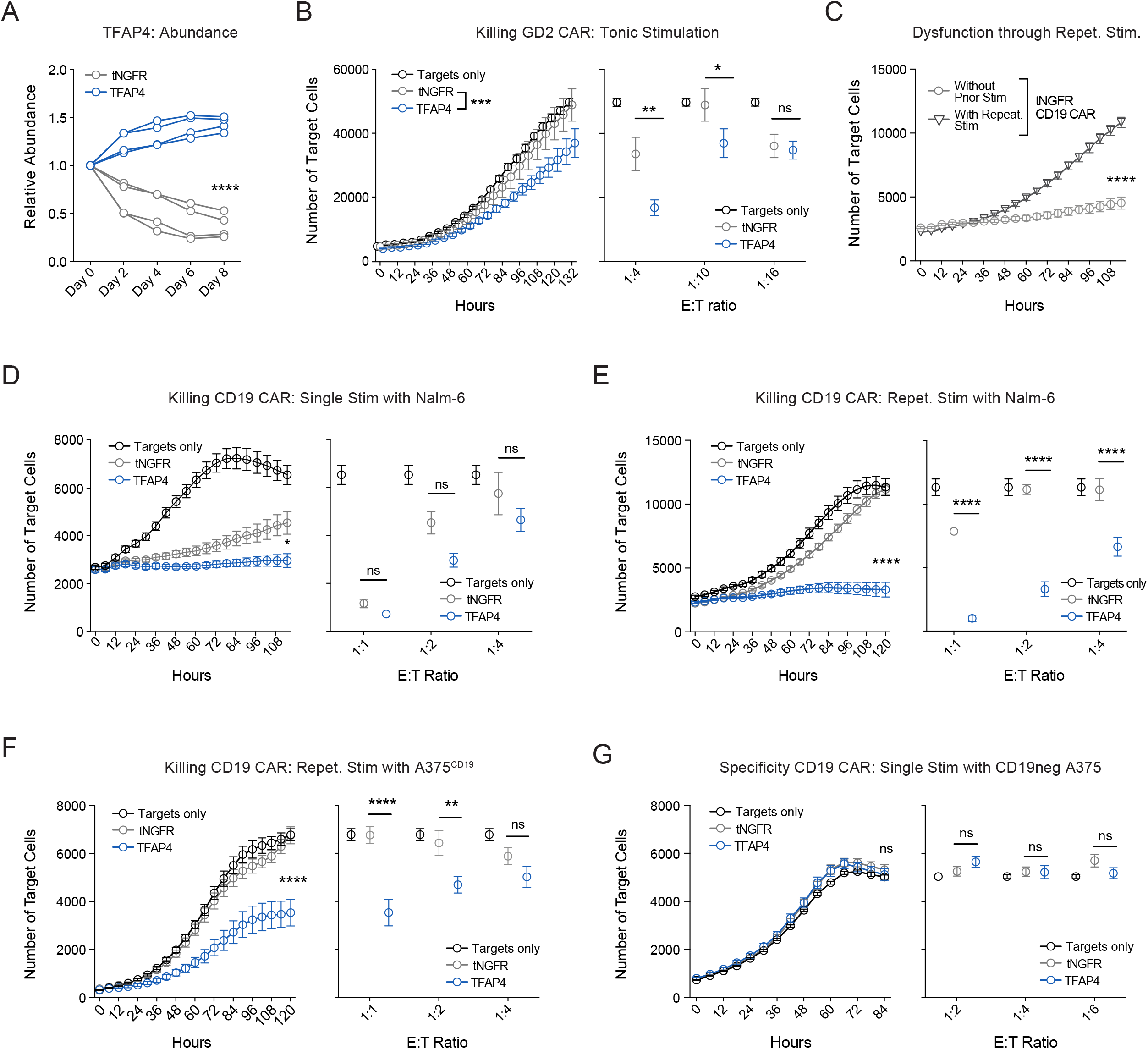
Extended TFAP4 Validation Data, Related to Figure 4. **(A)** The high affinity GD2 CAR was knocked into human T cells in combination with either TFAP4 or a control construct (tNGFR). Cells were sorted for CAR+/TCR-cells and TFAP4 and tNGFR T cells were co-cultured at a ∼50:50 ratio. Abundance of the NGFR+ cells was analyzed over time by flow cytometry. N = 2 individual donors in technical duplicates. Unpaired t test was used to calculate statistical significance on day 8. **(B)** tNGFR or TFAP4 GD2 CAR T cells were co-cultured with GD2+ target cells (Nalm-6/GFP/Luc/GD2) at various effector:target (E:T) ratios. Number of remaining target cells was calculated using the Incucyte system. N = 2 individual donors in technical triplicates. Two-way ANOVA was used to calculate statistical significance including Holm-Sidak’s multiple testing correction. Significance at last time point (132 hours) shown. Mean + SEM shown. Left panel shows 1:10 E:T ratio. **(C)** tNGFR CD19 CAR T cells were co-cultured with CD19+ target cells (Nalm-6) with or without prior 5x repetitive stimulation (CD19-positive A375s). Killing capacity of tNGFR CD19 CAR T cells was markedly decreased after going through repetitive stimulation. N = 2 individual donors in technical quadruplicates. Multiple unpaired t test was performed to determine statistical significance including Holm-Sidak’s multiple testing correction. Significance at last time point (114 hours) shown. E:T ratio = 1:2. Mean + SEM shown. **(D)** A CD19 CAR in combination with tNGFR or TFAP4 was knocked into human T cells and co-cultured with CD19+ target cells (Nalm-6) at various E:T ratios. Number of remaining target cells/cancer cell killing was analyzed using the Incucyte system across various E:T ratios. N = 2 individual donors in technical quadruplicates. Two-way ANOVA was used to calculate statistical significance including Holm-Sidak’s multiple testing correction. Significance at last time point (114 hours) shown. Mean + SEM shown. Left panel shows 1:2 E:T ratio. **(E)** T cells were generated as described in D, subjected to the repetitive stimulation assay (A375) and then co-cultured with CD19+ target cells one more time (Nalm-6). Again, CD19 CAR T cells with synthetic TFAP4 knockin were better able to control tumor cell growth. N = 2 individual donors in technical quadruplicates. Two-way ANOVA was used to calculate statistical significance including Holm-Sidak’s multiple testing correction. Significance at last time point (120 hours) shown. Mean + SEM shown. Left plot shows 1:2 E:T ratio. **(F)** T cells were generated as described in D, subjected to the repetitive stimulation assay and then co-cultured with CD19+ target cells one more time (stimulations and final co-culture were with adherent cell line A375 that was modified to express CD19). Again, CD19 CAR T cells with synthetic TFAP4 knockin were better able to control tumor cell growth. N = 2 individual donors in technical quadruplicates. Two-way ANOVA was used to calculate statistical significance including Holm-Sidak’s multiple testing correction. Significance at last time point (120 hours) shown. Mean + SEM shown. Left plot shows 1:1 E:T ratio. **(G)** tNGFR or TFAP4 CD19 CAR T cells were co-cultured with CD19-negative target cells (A375). No elevated unspecific killing of the TFAP4 compared to the tNGFR construct was observed. Multiple unpaired t test between TFAP4 and tNGFR was performed to determine statistical significance including Holm-Sidak’s multiple testing correction. Significance at last time point (84 hours) shown. Left plot shows 1:4 E:T ratio. N = 2 individual donors in technical triplicates. Mean + SEM shown. Only significance between TFAP4 vs tNGFR is shown in B and D-G.

**Supplementary Figure S10.**
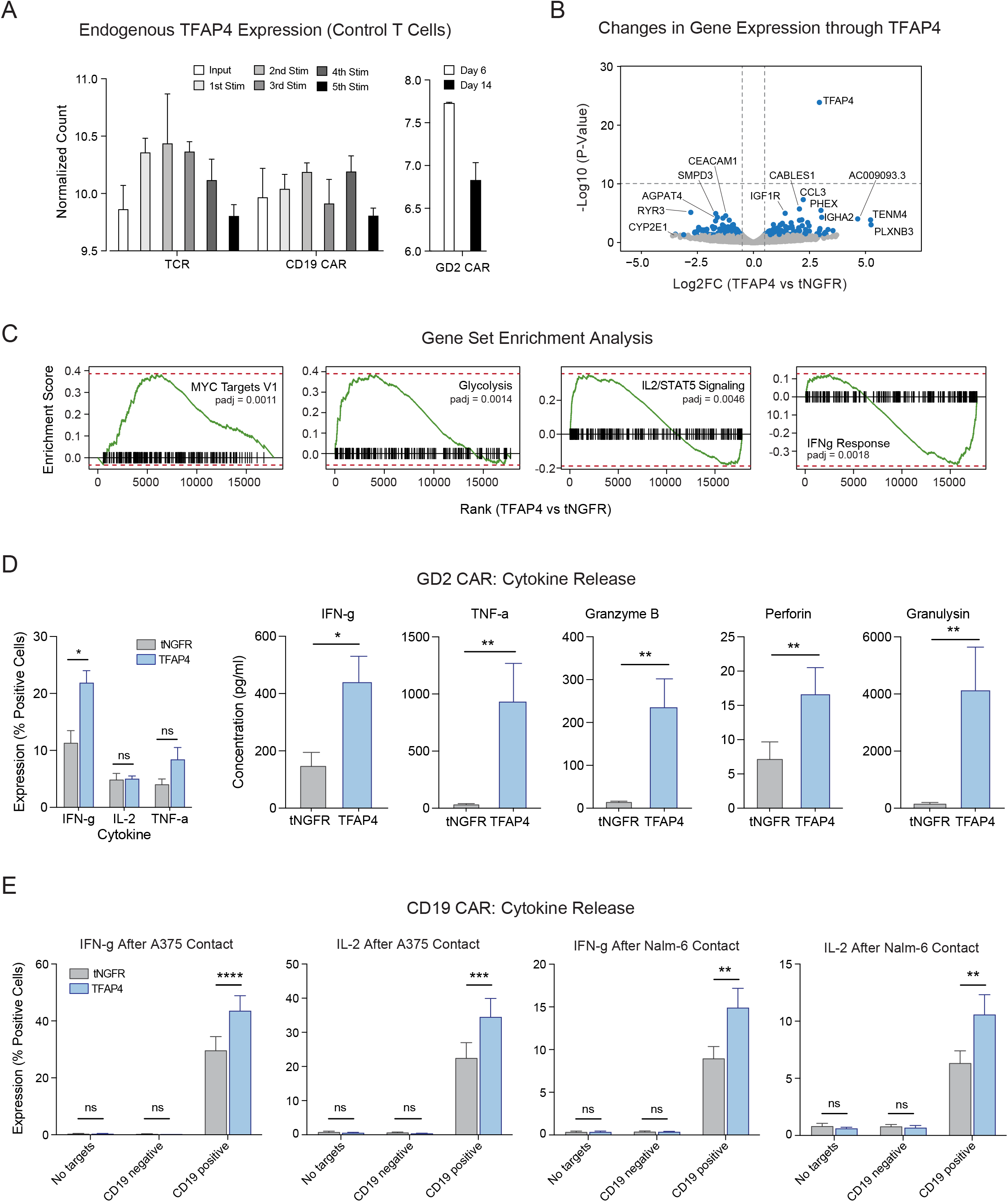
Extended TFAP4 Validation Data, Related to Figure 4. **(A)** Endogenous *TFAP4* expression was analyzed by bulk RNA-seq throughout the repetitive stimulation assay using the NY-ESO-1 TCR or CD19 CAR system and compared to endogenous *TFAP4* levels when culturing the GD2 CAR (tonic activation). While *TFAP4* expression peaked after the 2^nd^ stimulation in the TCR model, it was more heterogenous in the CD19 CAR model and decreased over time in the GD2 CAR model. N = 3 individual donors for the NY-ESO-1 TCR and CD19 CAR, n = 2 individual donors for the GD2 CAR. Mean + SEM shown. **(B)** RNA-seq was performed as described in Figure 4. Plot shows differentially expressed genes between TFAP4 and tNGFR KI GD2 CAR cells 7 days after electroporation. N = 2 individual donors. **(C)** Gene set enrichment analysis of RNA-seq data of GD2 CAR T cells showed that TFAP4 KI increased (relative to tNGFR KI) expression of MYC targets, glycolysis and IL2/STAT5 signaling genes and decreased expression of genes involved in the IFN-g response. N = 2 individual donors. **(D)** Cytokine production and secretion of TFAP4 KI vs tNGFR KI GD2 CAR T cells were analyzed after 24h co-culture with GD2+ target cells by intracellular cytokine stain (left panel) and Legendplex analysis of the supernatant (right panels). N = 2 individual donors in technical triplicates. Multiple paired t test was performed to determine statistical significance. Mean + SEM shown. **(E)** TFAP4 or tNGFR KI CD19 CAR T cells were co-cultured with either CD19 negative or CD19 positive target cells (A375 and Nalm-6). IFN-g and IL-2 production was evaluated by intracellular cytokine stain and confirmed elevated cytokine levels only in the presence of CD19-positive target cells. In the presence of CD19-negative target cells or in absence of target cells, TFAP4 KI CD19 CAR T cells did not release increased amounts of IFN-g and IL-2 compared to the control. Multiple paired t test was performed to determine statistical significance. N = 2 individual donors in technical triplicates. Mean + SEM shown.

**Supplementary Figure S11.**
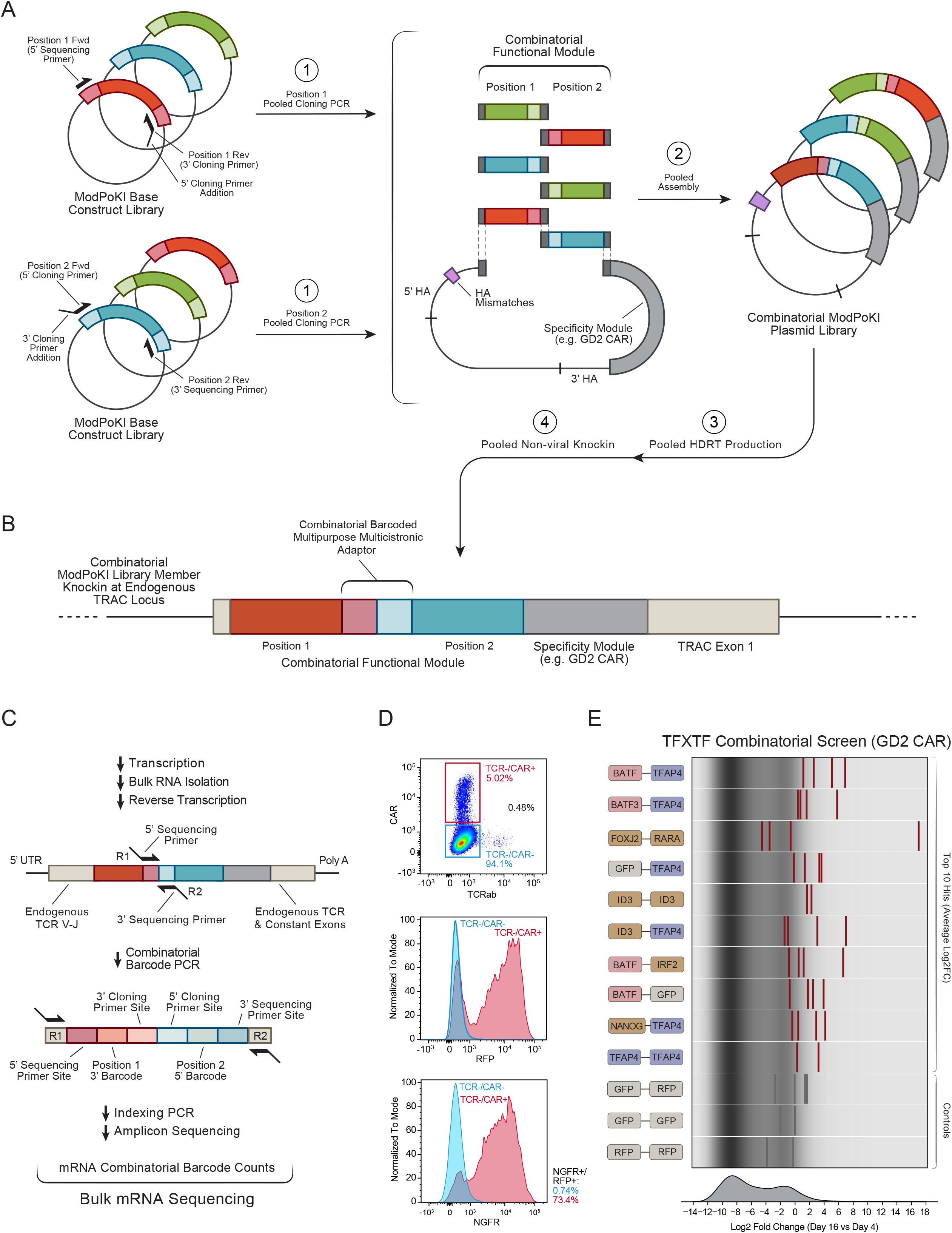
Combinatorial Knockin Strategy and Plasmid Representation of TFxTF Library, Related to Figure 5. **(A-B)** Schematic illustration showing the pooled assembly approach used to generate a combinatorial library of the tonic signaling GD2 CAR plus 100 transcription factors (TFs) x 100 TFs resulting in a ∼10,000-member library. The inserts for TF position 1 and 2 were separately generated by pooled PCRs off of the existing TF library. The backbone (consisting of the GD2 CAR plus homology arms) and the two inserts were assembled in a pooled Hifi DNA assembly reaction resulting in the combinatorial ModPoKI plasmid library. Double-stranded HDR template was generated by pooled PCR followed by non-viral pooled knockin into primary human T cells. **(C)** The resulting fusion region between the two transcription factors (combinatorial barcoded multicistronic adaptor) consisted of both the barcode of the insert in position 1 and the barcode of the insert in position 2. To read out the barcode region, mRNA was isolated, reverse transcribed into cDNA and amplified using the 5’ and 3’ sequencing adaptors to add Illumina Read 1 and Read 2. The indexing sequences were added in a second PCR step. Amplicon sequencing was performed and mRNA/cDNA combinatorial barcode counts were calculated. **(D)** Exemplary knockin of a control construct containing the GD2 CAR with tNGFR and RFP including the new combinatorial barcoded multicistronic adaptor sequences into primary human T cells. **(E)** The TFxTF combinatorial library was knocked into primary human T cells. As the GD2 CAR provides tonic signaling, T cells were cultured without addition of target cells. Cells were sorted on day 16 and day 4 after electroporation and the log2 fold change (log2FC) was calculated (day 16/day 4). Log2FC for the top 10 combinatorial TFxTF constructs is shown and compared to controls. N = 2 individual donors. Barcode representation of the two construct orientations x two donors is shown.

**Supplementary Figure S12.**
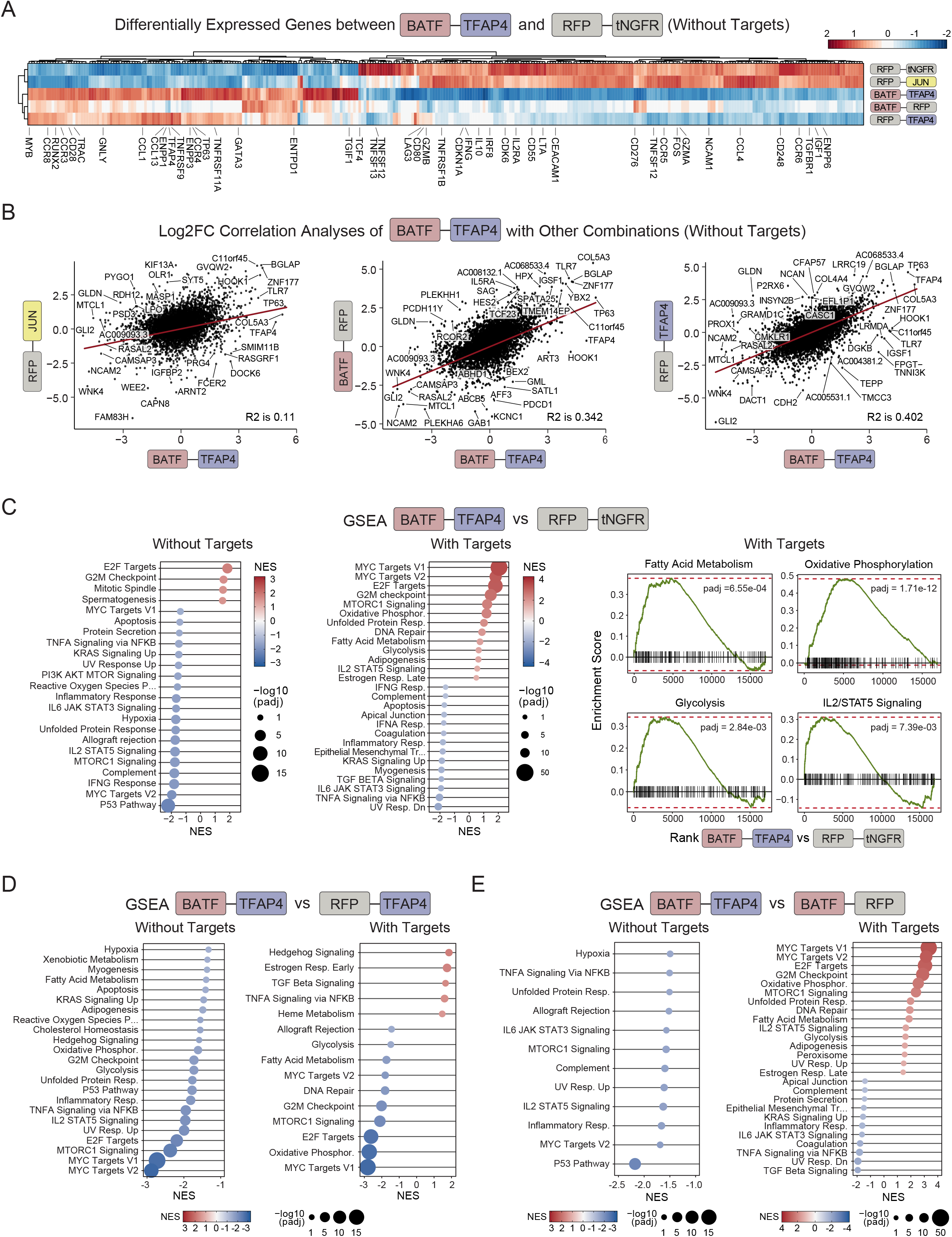
Transcriptomic Changes Driven by Combinatorial BATF-TFAP4 Knockin, Related to Figure 6. **(A)** Different combinatorial validation constructs were knocked into primary human T cells as described in Figure 6. Bulk RNA-seq was performed 14 days after electroporation. Differentially expressed genes between BATF-TFAP4 KI GD2 CAR T cells and RFP-tNGFR KI GD2 CAR T cells are plotted. N = 2 individual donors. **(B)** Log2 fold changes in gene expression between the tested KI condition and control KI (RFP-tNGFR) were correlated between BATF-TFAP4 and the other constructs. Correlation analyses indicated that BATF-TFAP4 KIs were most similar in gene expression with RFP-TFAP4 and BATF-RFP, while the correlation between BATF-TFAP4 KI cells and RFP-JUN KI cells was lower. N = 2 individual donors. Statistics were done using linear regression (lm function in R studio). **(C-E)** Gene set enrichment analysis of the BATF-TFAP4 combinatorial construct compared to RFP-tNGFR (C), RFP-TFAP4 (D) or BATF-RFP (E) without and with addition of GD2+ target cells for 24h is shown (day 14 vs day 15 after electroporation, respectively). Notably, after stimulation with target cells, expression of genes involved in fatty acid metabolism, glycolysis, oxidative phosphorylation and IL2/STAT5 signaling was increased in BATF-TFAP4 cells compared to RFP-tNGFR cells. N = 2 individual donors.

**Supplementary Figure S13.**
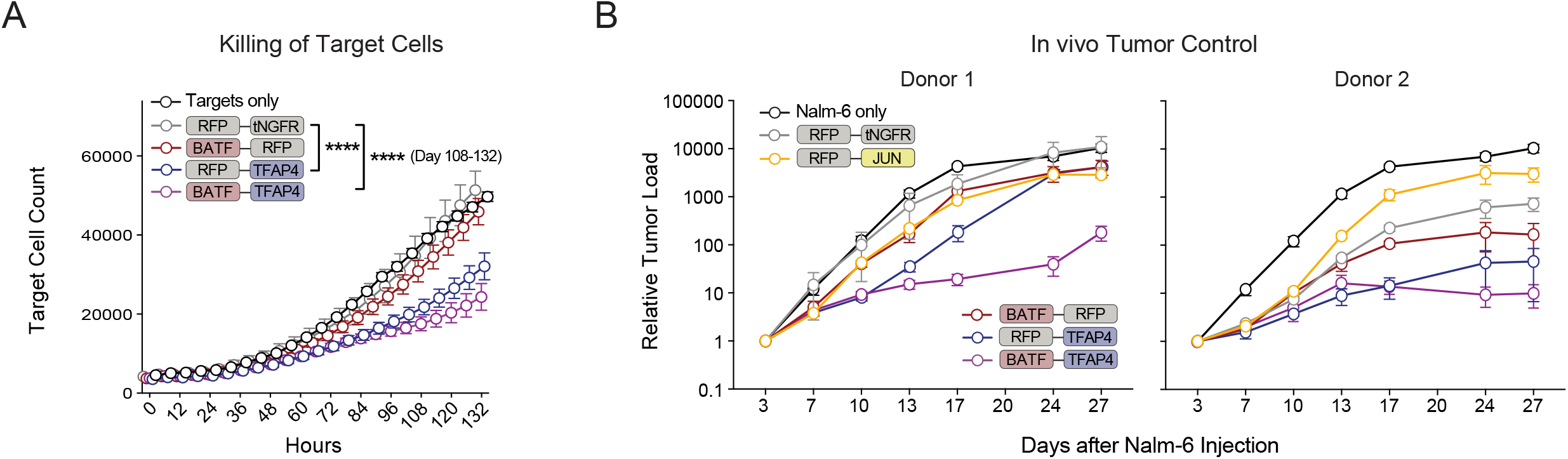
Functional Capacity of Combinatorial BATF-TFAP4 Knockin *in Vitro* and *in Vivo*, Related to Figure 6. **(A)** Combinatorial KI GD2 CAR T cells were co-cultured with Nalm-6/GFP/Luc/GD2 target cells and target cell killing was analyzed via Incucyte. BATF-TFAP4 combinatorial KI GD2 CAR T cells trended to outperform other combinatorial KI GD2 CAR T cells in terms of *in vitro* killing capacity. N = 2 individual donors in technical triplicates. 2-way ANOVA with Holm Sidak’s multiple comparisons test was performed to compare all constructs against the control (RFP-tNGFR). Mean+SEM shown. **(B)** Data from Figure 6G divided by donors. Mean + SEM shown.

## SUPPLEMENTARY TABLES

**Supplementary Table 1. Primer Sequences**

Cloning and sequencing primers used in this study

**Supplementary Table 2. Material**

Material used in this study

**Supplementary Table 3. TF Library DNA Sequences**

DNA sequences of the 100-member TF library including adaptor sequences and information on known functionality in T cells

**Supplementary Table 4. SR Library DNA Sequences**

DNA sequences of the 129-member SR library including adaptor sequences and information on switch receptor design

**Supplementary Table 5. TFxTF Library DNA Sequences**

DNA sequences of the combinatorial TFxTF library including adaptor sequences

**Supplementary Table 6. TF Library Screening Data**

Log2FC screening data of the TF library across TCR/CAR specificities and challenge modules

**Supplementary Table 7. SR Library Screening Data**

Log2FC screening data of the SR library across TCR/CAR specificities and challenge modules

**Supplementary Table 8. TFxTF Library Screening Data**

Log2FC screening data of the TFxTF library in the GD2 CAR tonic signaling screen

## METHODS

### Isolation and Culture of Primary Human T Cells

T cell isolation was done as previously described (Roth et al., 2020). Briefly, human T cells were isolated from leukapheresis products (Leukopaks, Stemcell, samples collected with approved Stemcell IRB) or TRIMA Apheresis (Blood Centers of the Pacific, San Francisco) using EasySep Human T Cell Isolation Kit (Stemcell). The use of human material is approved by the UCSF Committee on Human Research (CHR #13-11950). T cells were cultured in X-VIVO 15 media (Lonza Bioscience) supplemented with 5% fetal bovine serum (FBS), 50 µM 2mercaptoethanol (Thermo Fisher Scientific), and 10 mM N-Acetyl-L-Cysteine (VWR). Prior to electroporation, T cells were stimulated for 48 hours at 1e6 cells per ml of media containing 500 U/ml IL-2 (R&D Systems), 5 ng/ml IL-7 (R&D Systems), 5 ng/ml IL-15 (R&D Systems), and CTS (Cell Therapy Systems) CD3/CD28 Dynabeads (Thermo Fisher Scientific, bead:cell ratio 1:1). After nucleofection, T cells were cultured in X-VIVO 15 media containing 500 U/ml IL-2 unless otherwise stated and split every two to four days.

### Generation of Plasmid Libraries for Pooled Knockin

The 231 constructs included in the pooled knockin library (Supplementary Tables 3 and 4) were designed using the Twist Bioscience codon optimization tool and were commercially synthesized and cloned (Twist Bioscience) into a custom pUC19 plasmid containing the NY-ESO-1 TCR sequence (except for HIF1A, JUN and TCF7 constructs that were cloned individually using gBlocks Gene Fragments (Integrated DNA Technologies)). Individual pooled plasmid libraries were created by pooling single construct plasmids into respective libraries (transcription factors, 100 members; surface receptor constructs, 129 members; controls, 2) or in one complete pool. The CD19 CAR plasmid pool was created in a pooled assembly fashion by amplifying constructs from the TCR plasmid pool as a DNA template. PCR amplification (Kapa Hot Start polymerase, Roche) produced a pooled library of amplicons with small overhangs homologous to a pUC19 plasmid containing the CD19 CAR HDR sequence. The amplicon pool was treated with DpnI restriction enzyme (New England Biolabs, NEB) to remove residual circular TCR plasmids, bead purified (Sera-Mag SpeedBeads), and eluted into H2O. We then used Gibson Assembly (NEB) to construct a plasmid pool containing all 231 library members and knockin controls, plus the new CAR sequence. The CD19 CAR plasmid pool was bead purified, transformed into Endura electrocompetent cells (Lucigen) and maxiprepped (Plasmid Plus Midi or Maxi Kit, Qiagen) for further use. The GD2 CAR libraries were generated in a similar way. While the NY-ESO-1 TCR libraries were pooled at the plasmid stage (plasmids were synthesized individually), all other plasmid libraries in this project (anti-CD19 CAR, anti-GD2 CAR, combinatorial library) were generated by pooled Gibson assembly of the plasmids. The CD19 CAR sequence used in this study was kindly provided by Tobias Feuchtinger, Dr. von Hauner Children’s Hospital, University Hospital, LMU Munich. The GD2 CAR sequence was kindly provided by Crystal Mackall and Robbie Majzner, Stanford (Lynn et al., 2019). Primer sequences are listed in Supplementary Table 1.

### Generation of Combinatorial Libraries for Pooled Knockin

The combinatorial GD2 CAR plasmid libraries were generated by pooled Gibson assembly of a GD2 CAR pUC19 backbone as well as TF insert 1 and TF insert 2. The backbone included the published GD2 CAR sequence (Lynn et al., 2019) with CD28 co-stimulation and mutations in the IgG1 CH2 region to increase tonic signaling (Watanabe et al., 2016) (kindly provided by Crystal Mackall and Robbie Majzner as described above). The inserts were PCR-amplified out of the pre-existing TF library using primers that removed the 5’ barcode of the first insert and the 3’ barcode of the second insert and added a constant linker in between the two combinatorial inserts. The GD2 CAR backbone, the pools of insert 1 and the pools of insert 2 were assembled using NEBuilder HiFi DNA Assembly Master Mix (NEB). The assembled product was bead-purified using Sera-Mag SpeedBeads (Thermo Fisher Scientific), transformed into Endura electrocompetent cells (Lucigen) and midi- or maxiprepped (Plasmid Plus Midi or Maxi Kit, Qiagen) for further use. Primer sequences are listed in Supplementary Table 1.

### Homology Directed Repair Template (HDRT) Generation

HDRTs were produced as previously described (Roth et al., 2020). In brief, TCR or CAR plasmid pools were used as templates for PCR amplification (KAPA HiFi HotStart ReadyMix, Roche) to generate double-stranded DNA templates including truncated Cas9 target sequences (Nguyen et al., 2020). Templates were bead-purified as described above and eluted into H2O. The concentrations of eluted HDRTs were normalized to 500-1,000 ng/µL. HDRT amplification was confirmed by gel electrophoresis in a 1.0% agarose gel. The templates for arrayed knockin of the different single constructs or combinations during the validation stage were generated in a similar way. Instead of libraries, single constructs served as templates for the PCRs. In all cases, primers were used that added a truncated Cas9 target sequence (Nguyen et al., 2020).

### Cas9 RNP Electroporation

Electroporation was done as previously described (Roth et al., 2020). To produce ribonucleoproteins (RNPs), crRNA and tracrRNA (stock 160µM, both Dharmacon) were mixed 1:1 by volume, and annealed by incubation at 37C for 30 min to form an 80 µM guide RNA (gRNA) solution. Poly-L-glutamic acid (PGA, stock 125mg/ml, Sigma) was mixed with gRNA at 0.8:1 volume ratio prior to complexing with Cas9-NLS (QB3 Macrolab) for final volume ratio gRNA:PGA:Cas9 of 1:0.8:1 (Nguyen et al., 2020). These were incubated at 37C for 15 min to form a 14.3µM RNP solution. RNPs and HDRTs were mixed with T cells before electroporation (3.5µl of RNP with 500ng-1µg=1µL of HDRT). Bulk T cells were resuspended in electroporation buffer P3 (Lonza Bioscience) at 0.75e6 cells per 20µl (per well) and transferred to a 96-well electroporation plate together with 4.5µL of RNP/HDRT mix per well. Pulse code EH115 was used on a 4D-Nucleofector 96-well Unit (Lonza Bioscience). Cells were rescued in X-VIVO 15 without cytokines for 15min and then cultured in X-VIVO 15 with 500 U/ml IL-2.

### Flow Cytometry and FACS

For flow cytometric analysis, T cells were centrifuged at 300g for 5 min and resuspended in flow buffer (PBS/2%FCS) containing the respective antibody mix (see Supplementary Table 2). For NY-ESO-1 TCR constructs, cells were stained for 12 min with Dextramer-HLA-A*0201/SLLMWITQV-PE (Immudex) before adding surface antibodies. For GD2 CAR constructs, cells were stained for 15 min at 4C with Alexa Fluor 647 AffiniPure F(ab’)2 Fragment Goat Anti-Mouse IgG, F(ab’)2 fragment specific (Jackson ImmunoResearch), washed once with flow buffer (PBS with 2mM EDTA), resuspended in 100ul 2% mouse serum in PBS, incubated for 10 min at 4C, and washed again before surface stain antibodies were added. After another 10 min incubation, cells were washed again and resuspended in wash buffer, then analyzed on an Attune NxT Flow Cytometer (Thermo Fisher Scientific) or BD LSRFortessa (BD Biosciences). For CD19 CAR constructs, detection through the integrated myc tag was done according to the manufacturer’s instructions (Myc-Tag (9B11) Mouse mAb (Alexa Fluor 647 Conjugate), Cell Signaling Technology).

### Single Stimulation Screens

One day prior to set-up of the screen, 2.5e6 A375s were plated per T75 flask in complete RPMI media (RPMI plus NEAA, Glutamine, Hepes, Pen/Strep, sodium pyruvate (all Thermo Fisher Scientific) and 10% FCS (Sigma-Aldrich, St. Louis, Missouri, USA)) assuming that they double within 24 hours. One day later (= seven days after electroporation), edited T cell pools were counted and washed once. 10e6 T cells were transferred to TRI Reagent (Sigma-Aldrich) representing the input population for amplicon sequencing. 10e6 T cells per screening condition were transferred to one T75 flask in 20 ml of X-VIVO 15 (Lonza Bioscience) supplemented with 5% FCS, 2-Mercaptoethanol (Thermo Fisher Scientific), N-Acetyl-L-Cysteine (VWR) and 50 U/ml IL-2 (Proleukin). For A375 conditions, cRPMI was removed and flasks were filled up with 20 ml of X-VIVO 15 plus additives and 10e6 T cells. For Nalm-6 conditions, 5e6 Nalm-6 cells were added per T75 flask. In the stimulation conditions, T cells were stimulated with Dynabeads CD3/CD28 CTS (Thermo Fisher Scientific) at a 1:1 bead: cell ratio (“signal 1+2 stim”) or a 5:1 ratio (“signal 1+2 excess stim”). For CD3 stimulation only (“signal 1 stim” condition), T cells were incubated with NY-ESO-1 specific dextramer (Immudex) for 12 min at RT (1:50 dilution), washed once and transferred to a T75 flasks. After two days, 10 ml of X-VIVO 15 were added to all conditions including supplements and 50 U/ml IL-2. Another two days later, cells were counted and 10e6 cells were transferred to TRI Reagent (Sigma-Aldrich) for RNA isolation and amplicon sequencing. The Nalm-6 cell line used in the TCR single stimulation screens had been previously modified to express the NY-ESO-1 antigen on HLA-A2 (in addition to GFP/Luc) and was a kind gift from Justin Eyquem, Gladstone-UCSF Institute of Genomic Immunology, San Francisco.

### Repetitive Stimulation and Tonic Signaling Screens

One day prior to the start of the repetitive stimulation screen, A375 cells were counted and transferred to 24-well plates (50,000 cells per well in 1 ml of complete RPMI media) assuming that they double within 24 hours. One day later, edited T cell pools were counted and 10e6 cells were frozen in TRI Reagent (Sigma-Aldrich) for amplicon sequencing (input population). Media of the A375 cells was removed. 100,000 edited T cells (NY-ESO-1 multimer or CAR positive, ∼1:1 effector:target ratio) were transferred to each well of the 24-well plate and co-cultured with the A375 cells in 2 ml of X-VIVO 15 containing supplements plus 50 U/ml IL-2. 24 hours later, fresh A375 cells were plated as described above. One day later, media of the new A375 plate was removed and replaced by 1 ml of fresh X-VIVO 15 plus 1 ml of the T cell suspension from the first plate including 50 U/ml IL-2 calculated on the total volume per well. The rest of the T cells were counted and 10e6 cells were transferred to TRI Reagent (Sigma-Aldrich) for amplicon sequencing. The procedure was repeated every other day for a total number of five stimulations with target cells. Multiple wells of the 24-well plates were used per screen to reach cell coverage. For tonic signaling screens, the GD2 CAR libraries were knocked into T cells. GD2 CAR T cells were not stimulated with target cells as the GD2 CAR is known to drive tonic activation. Cells were harvested on day 4, 8, 12 and 16 after electroporation and transferred to TRI Reagent (Sigma-Aldrich). Combinatorial tonic signaling screens were performed in a similar way (harvest day 4 and day 16). When working with CD19 CARs in combination with A375 cells, we used CD19 overexpressing A375 cells (SFFV promoter knocked in upstream of endogenous CD19).

### Barcode/Amplicon Sequencing

Genomic DNA (pilots) or RNA (unless otherwise noted) was isolated from input and output population using Quick-DNA and Direct-zol RNA kits, respectively (Zymo Research). RNA was reverse transcribed into cDNA using Maxima H Minus Reverse Transcriptase (Thermo Fisher Scientific). The sequencing library was generated by two PCRs. PCR1 was performed using KAPA HiFi HotStart ReadyMix (Roche) for 18 cycles. Amplicons from PCR1 were bead-purified. For PCR2, NEB Next Ultra II Q5 polymerase (NEB) was used for 10 cycles to append P5 and P7 Illumina sequencing adaptors. The PCR2 product was bead-purified, normalized libraries were pooled across samples and sequenced on a MiniSeq or NextSeq500 (Illumina). Barcode distribution was analyzed and log2 fold change of barcode representation in output vs input population was calculated to detect changes in abundance. Primer sequences are shown in Supplementary Table 1.

### Competition Assay

For validations, after arrayed knockin of the different constructs, cells were sorted and competition assay was set up on day 8-10 after electroporation. T cells were cultured at a ∼50/50 ratio with control T cells in X-VIVO 15 containing 50 U/ml IL-2. The cell ratio was confirmed by flow analysis of the cell mixes and exact percentage of control T cells was determined at baseline level (NGFR expression). Changes in cell ratio were normalized based on percentages on day 0 of the assay.

### Activation Marker and Phenotype Analysis

For GD2 CAR validation assays, activation marker expression (CD25, CD69) was analyzed by flow cytometry on day 8 after electroporation. CD62L/CD45RA expression levels were analyzed by flow cytometry 14 days after electroporation.

### RNA-Sequencing (Bulk RNA-Seq)

For control TCR RNA-seq experiments (Figure S5C-E), a dataset from a manuscript recently accepted for publication was analyzed ((Carnevale et al., 2022), GSE204862). For GD2 CAR validation experiments, edited cells were sorted for CAR+/TCR-expression on day 7 (single inserts) or on day 6 and day 14 (combo inserts) after electroporation. On day 14, one part of the sorted population was stored in TRI Reagent for RNA-seq (Sigma-Aldrich), the other part was stimulated with Nalm-6/GFP/Luc/GD2 cells at a 1:1 E:T ratio. After 24h, the stimulated T cells were sorted again for CAR+/TCR-cells and stored in TRI Reagent (Sigma-Aldrich). RNA was isolated using Direct-zol RNA kits (Zymo Research). RNA was prepared for sequencing as previously described (Cortez et al., 2020) by the Functional Genomics Laboratory at UC Berkeley and sequenced by the Vincent J. Coates Genomics Sequencing Laboratory at UC Berkeley. Kallisto was used to map the reads to the human reference transcriptome and genes with zero counts in more than 80% of samples were removed from the analysis. DESeq2 R package was used for differential gene expression, fgsea package for gene set enrichment analysis (GSEA) with MSigDB v7.2 hallmark gene sets as reference gene lists. Nalm-6/GFP/Luc/GD2 were a kind gift from the Mackall lab (Stanford) and were reported to have an STR profile that was a ∼60% match to Nalm-6, suggesting some degree of mutation.

### Modular Pooled Knockin Screening with Single-Cell RNA Sequencing (ModPoKI-Seq)

PoKI-Seq was performed as previously published (Roth et al., 2020). Since the knockin barcodes were closer to the 5’ end of the transcript compared to the previous PoKI design, Chromium Single Cell 5’ Reagent Kit, v1 chemistry (10x Genomics) was used according to the manufacturer’s protocol. NY-ESO-1 TCR-positive cells were sorted by FACS, counted and resuspended at 1,000 cells/µL in PBS with 1% FCS. After GEM (Gel Bead-In Emulsions) recovery, the mRNA library was converted to cDNA, amplified for 11 cycles, and quantified with Agilent Bioanalyzer High Sensitivity. 75% of the amplified cDNA material was carried through for transcriptome library preparation according to the manufacturer’s protocol. The remaining 25% of amplified cDNA was used for amplicon sequencing of the knockin barcodes. The cDNA was enriched for knockin barcodes using a nested PCR strategy with Kapa HiFi HotStart Ready Mix (Roche) for 8 cycles per round. For the first PCR, 0.5µM each of ModPoKI_Seq_1_forw primer and ModPoKI_Seq_1_rev primer was used. Amplified products were purified with 0.8x SPRIselect Reagent Kit (Beckman Coulter) and eluted in 10µL nuclease-free water. The libraries were further enriched with a second PCR using 0.5µM each of ModPoKI_Seq_2_forw primer and ModPoKI_Seq_2_rev primer. Amplified products were purified with 0.8x SPRIselect Reagent Kit (Beckman Coulter) and eluted in 15µL nuclease-free water. Lastly, index PCR was performed with Kapa HiFi HotStart Ready Mix (Roche) for 8 cycles with 2.5µL each Nextera Chromium i7 Sample Indices N Set A (PN 3000262) and 0.5µM ModPoKI_Seq_index primer. Amplified products were purified with 0.8x SPRIselect Reagent Kit (Beckman Coulter) and eluted in 45µL nuclease-free water. Samples were pooled and sequenced on a NovaSeq S4 flow cell with 20% PhiX using read parameters 30×8×98. Fastq files were mapped to the human transcriptome (10x Genomics Cell Ranger, v5.0.0) and a custom knockin barcode reference and analyzed using Seurat (v4.1.1) (Butler et al., 2018). A small fraction (<0.4%) of A375 target cells forming a distinct cluster were removed from the dataset after manual inspection

### *In Vitro* Intracellular Cytokine Assay and Legendplex

T cells were stimulated with target cells at a 1:1 E:T ratio for 24h. 1x Brefeldin A (Thermo Fisher Scientific) was added to the culture for 4 hours. Cells were spun down and supernatant was frozen for Legendplex analysis (LEGENDplex Human CD8/NK Panel 13-plex, BioLegend, according to the manufacturer’s information). Cells were stained for surface markers and intracellular cytokines (see Supplementary Table 2) using the FIX & PERM Cell Fixation & Cell Permeabilization Kit (Thermo Fisher Scientific) according to the manufacturer’s information. For GD2 CAR assays, Nalm-6/GFP/Luc/GD2 were used as target cells (kind gift from the Mackall Lab, as described above). For CD19 CAR assays, A375s with CD19 (SFFV promoter knocked in upstream of endogenous CD19) and without CD19 expression (WT) or Nalm-6 cells with and without CD19 expression (CD19 knockout, kind gift from the Eyquem Lab, as described above) were used.

### TOX Stain

Intracellular transcription factor stains were done using the eBioscience Foxp3/ Transcription Factor Staining Buffer Set (Thermo Fisher Scientific) according to the supplier’s information. A list of flow antibodies is provided in Supplementary Table 2.

### *In Vitro* Killing Assay

For Incucyte assays with Nalm-6/GFP/Luc/GD2 (gift from the Mackall Lab, as described above), flat bottom 96-well plates were coated with 50µl of 0.01% poly-L-ornithine (PLO) solution (Sigma) for 1 hour. PLO was removed and plates were dried for 30-60 min. 10,000 Nalm-6/GFP/Luc/GD2 per well were mixed with sorted T cells in various effector:target (E:T) ratios. For Incucyte assays with A375 target cells (RFP+), 1,500 A375 cells were plated into flat bottom 96-well plates 24h before start of the assay. T cells were added in various E:T ratios one day later (assuming that the A375 cells doubled within 24h). The assay media consisted of X-VIVO 15 as described above, supplemented with 500 U/ml IL-2 and 1X Glucose Solution (Thermo Fisher Scientific). Cell counts were analyzed every six hours using the Incucyte Live Cell Analysis System (Essen BioScience). When working with CD19 CARs in combination with A375 cells, we used CD19 overexpressing A375 cells (SFFV promoter knocked in upstream of endogenous CD19).

### *In Vivo* Mouse Model

NOD/SCID/IL2Rg-null (NSG) mice were purchased from Jackson Laboratory. 8-12 weeks old female mice were used and mouse experiments were performed under an approved UCSF Institutional Animal Care and Use Committee protocol. For tumor control and survival analyses, mice were injected IV with Nalm-6/GFP/Luc/GD2 cells (gift from the Mackall Lab, as described above) on day 0. Three days later, edited human T cells were injected IV (T cell count was calculated based on CAR+ T cells). Nalm-6 and T cell doses are indicated in the figure legends. T cells were TCR-depleted one day before injection using EasySep Human TCR Alpha/Beta Depletion Kit (Stemcell) to avoid Graft-versus-Host disease in the mice by unedited cells. Knockin rates were adjusted between groups by adding TCR-negative T cells without CAR knockin right before injection. These cells were generated simultaneously with the therapeutic cells from the same donor and treated the same way except no HDR template was added during electroporation. For imaging, 200 µL (3 mg) of D-Luciferin Potassium Salt (Gold BioTechnology) were injected IP and mice were imaged using an IVIS Spectrum *In Vivo* Imaging System (PerkinElmer) once/twice per week.

### Statistics

Statistical details for all experiments can be found in the figure legends. Ns = not significant, * <0.05, **<0.01, ***<0.001, ****<0.0001.

